# From populations to pan-genomes: investigating the role of ecology and genomic architecture in maintaining species boundaries in the porcini mushroom, *Boletus edulis*

**DOI:** 10.1101/2023.09.05.556370

**Authors:** Keaton Tremble, Etienne Brejon Lamartinière, Alexander J. Bradshaw, Roberto Flores Arzú, Rytas Vilgalys, Joseph Hoffman, Bryn T.M. Dentinger

## Abstract

Hybridization among diverged species is common across all domains of eukaryotic life, but appears to be particularly common in fungi. However, some fungi exhibit a greater tendency for hybridization than others and it is unclear what mechanisms facilitate or prevent hybridization. Here, we generated 253 whole genome sequences and 22 reference genomes within the globally distributed ectomycorrhizal species complex *Boletus edulis* Bull. and use a multi-faceted genomic approach to identify patterns of ongoing hybridization and determine whether hybridization is mediated by changes in genome structure, ecology, or both. We found that hybridization is common among species of *B. edulis* despite 2.1 million years of divergence. However, not all lineages that hybridize today exhibit patterns of introgression, indicating the presence of strong reproductive boundaries among some species. Using a pan-genomic approach we find that genome structural variation is abundant within *B. edulis*, but that the presence of ongoing introgression does not correlate with genome structural similarity or overall gene content. Instead, we find that the composition of ecologically relevant gene families more accurately predicts the presence of introgression among lineages and *B. edulis* as a group may specialize on chitin decomposition. Altogether, we show that ecological preferences are likely the primary driver of reproductive barriers in *B. edulis*.

## 1 Introduction

Hybridization events carry enormous evolutionary potential as drivers of speciation, extinction, and adaptive radiations [1]. However, despite their evolutionary importance, the fundamental mechanisms that govern the hybridization process are poorly understood across eukaryotes. While interspecific hybridization occurs across all life, fungi exhibit an exceptional capacity among eukaryotes for genetic exchange among distantly related groups [2]. For example, species of the genus *Saccharomyces*, commonly known as the yeasts, have long been shown to readily hybridize, even among parental lineages with an average level of protein divergence roughly equivalent to that of humans and chickens [3]. Moreover, hybridization events in Fungi appear to result in genetic exchange among species, termed introgression, at an exceptional rate [4, 5] which often drives adaptation to novel niches [2, 6]. It is undeniable that hybridization plays a dynamic role in fungal evolution. However, we lack a fundamental understanding of why hybridization is so common in fungi, and what are the mechanisms that mediate this phenomenon; why do certain fungi hybridize, while others do not?

The mechanisms that dictate successful hybridization and introgression have been studied in depth in animals and some plants, but have received less attention in fungi where they may play a reduced role. For many animals [7, 8], and some plants [9], pre-mating mechanisms (mechanisms that prevent mating among individuals) such as assortative (non-random) mating and allopatric isolation are often sufficient to reinforce species boundaries, preventing hybridization. However, assortative mating may be largely uncommon in the Fungi, and evidence that suggests assortative mating reinforces species boundaries is sporadic [10, 11, 12]. Furthermore, allopatric isolation may be less of an insurmountable barrier in fungi, as they often possess far greater dispersal capacity than animals or plants [13, 14, 15]. Thus, post-mating reproductive barriers (mechanisms that act after mating among individuals), are likely the most common mechanisms that mediate hybridization [2, 16]. There are numerous post-mating mechanisms that can prevent hybridization and introgression in fungi [2], yet, these mechanisms mediate hybridization through more or less two overarching categories; 1) reducing genomic compatibility of parents, or 2) reducing ecological fitness of offspring. While these overarching categories have been well investigated in laboratory settings in a select few model taxa, they have rarely if ever been investigated in natural populations where they may be ineffectual at large scales. Thus, it is uncertain whether genomic and/or ecological barriers are necessary to prevent fungal hybridization and introgression.

Genomic structural variation is perhaps the most common mechanism that interferes with genomic compatibility of hybridizing parents in fungi [17, 18] and plays an integral role in mediating hybridization when large chromosomal inversions, translocations, insertions, and deletions lead to hydrid dysfunction. This occurs when rearrangements result in the incorrect recombination of chromosomes and the formation of underdominant gametes [19, 20]. However, this common postzygotic mechanism may not be an insurmountable barrier in fungi. For example, it has been demonstrated that hybridization of heteroploid parental species can lead to sexually viable aneuploid hybrids [21, 22], and that the distinct lack of intraspecific genomic synteny, termed mesosyteny, is a diagnosable characteristic of both sexual and asexual *Dothideomycetes* [23]. This suggests that structural variation, which would be expected to produce hybrid dysfunction, is less consequential for hybridization in *Fungi* than in other eukaryotes. In contrast, structural variation is the main cause of hybrid sterility in other fungi. After hybridization of two *Saccharomyces* species, chromosome pairs failed to segregate during meiosis in 40% of all sporulation events [24]. In another study, Yadev et al. (2020) induced substantial inter-chromosomal rearrangements in *Cryptococcus neoformans*, and found that only 3% of isolate pairings produced viable offspring [25]. These examples clearly show that at least some fungi are not immune to the impacts of genomic structural variation and are not entirely unique from other eukaryotes. It is therefore unclear to what extent large structural variations mediate the outcome of hybridization in fungi.

The role that ecology plays in mediating hybridization has received substantially less attention than genomic compatibility, although it could conceivably play an equally important role [26]. Parental lineages that exhibit distinct ecological lifestyles may be more likely to produce maladapted hybrids. This would lead to strong barriers to hybridization among parents with low ecological similarity. Yet, without niche differentiation from their parents, new hybrid genotypes can be overcome by competition and/or gene flow from parental populations [27, 26]. As evidence, hybrid lineages often exhibit dramatic niche shifts [2] and are ecologically distinct from their parents [28]. This may suggest that parents with low ecological similarity are more likely to produce hybrids that survive. Regardless, the role that ecology plays in mediating hybridization has almost exclusively been viewed from the lens of hybrid fitness, and not parental lineage ecological similarity [29, 26].

The prized edible porcini mushroom, *Boletus edulis*, is a globally distributed ectomycorrhizal species complex with an exceptional ecological amplitude. *B. edulis* can be found in nearly every temperate ecosystem in the northern hemisphere, from the tropics in Guatemala, to north of the arctic circle in Alaska [30]. Tremble et al. (2022) conducted an initial phylo- and population genomic survey of the complex and found that *B. edulis* segregates into six phylogenetic species that have diverged over the past 2.6 million years. These phylogenetic species (here called lineages for simplicity) exhibit cryptic but geographically distinct population structure that does not correspond with previously identified ecological or morphological forms. Importantly, they found that some lineages of *B. edulis*, irrespective of geographic proximity or periodic allopatry, exhibited strong signatures of hybridization while others showed little to none [13]. The expansive geographic distribution, ecological and cultural importance, and contrasting patterns of hybridization makes *B. edulis* an ideal system to investigate the mechanisms that mediate hybridization among long-diverged organisms.

To identify whether genomic similarity, and/or ecological similarity, prevent or allow for hybridization among long diverged groups we conducted perhaps the largest genomic survey of a non-model eukaryotic species complex to date. To this end, we undertook four main tasks: 1) generate substantial new genomic resources across the *B. edulis* complex, including 23 reference genome assemblies and 220 whole genome sequences, 2) identify signatures of gene flow among long-diverged lineages, 3) characterize the degree of genomic structural variation and synteny among lineages, and 4) identify signatures of ecological differentiation among lineages of *B. edulis* by testing for changes in abundance of ecologically relevant gene families. We investigated whether the presence/absence of hybridization and introgression among lineages correlates with genomic and/or ecological similarity. We hypothesized that both structural similarity and ecological similarity are strong mediators of hybridization. We demonstrate that in natural populations of fungi, hybridization and introgression occurs irrespective of genomic architectural similarity and may be strongly mediated by ecological similarity.

## 2 Results

### 2.1 Phylogenetic analysis of cyto-nuclear discordance reveal patterns of hybridization among lineages of *B. edulis*

To identify patterns of hybridization among lineages of *B. edulis*, we used a summary coalescent phylogeneomic approach to compare the nuclear and mitochondrial genomes of 253 specimens (Table S1) and identify clear patterns of cyto-nuclear discordance indicative of hybridization. Cophylogenetic analysis of the *B. edulis* nuclear genome and mitochondrial genome reveled substantial cyto-nuclear discordance (Fig. 1A. see supplemental information for full results). Three lineages that appear monophyletic in the nuclear species tree Alaska-Siberia (AK), West Coast North America (WC), and Colorado (CO) were polyphyletic in the mitochondrial tree, with well-supported nodes leading to separated clusters. Empirical tests for signatures of hybridization within the mitochondrial genome using the HyDe algorithm suggest that these patterns of discordance are likely the result of introgression and not solely incomplete lineage sorting (ILS) (Table S4, see supplementary results for additional details). Further, we identified at least six individuals with hybrid genotypes (Fig. 1 arrows on right); where, an individual was placed within the monophyletic lineage consistent with its collection location in the nuclear species tree, but was placed within another lineage in the mitochondrial tree with significant nodal support (BS > 0.8). Two individuals from the AK lineage were placed in the European (EU) mitochondrial group, two individuals from British Columbian (BC) lineage in the WC lineage, one from the East Coast North America (EC) lineage in the AK lineage, and one from the WC lineage was placed within the BC lineage. Moreover, these hybrids also exhibited significant signatures of genetic introgression within their mitochondiral genome (Zscore > 2, P < 0.05, Table S5), again exceeding that expected for ILS.

**Figure 1:**
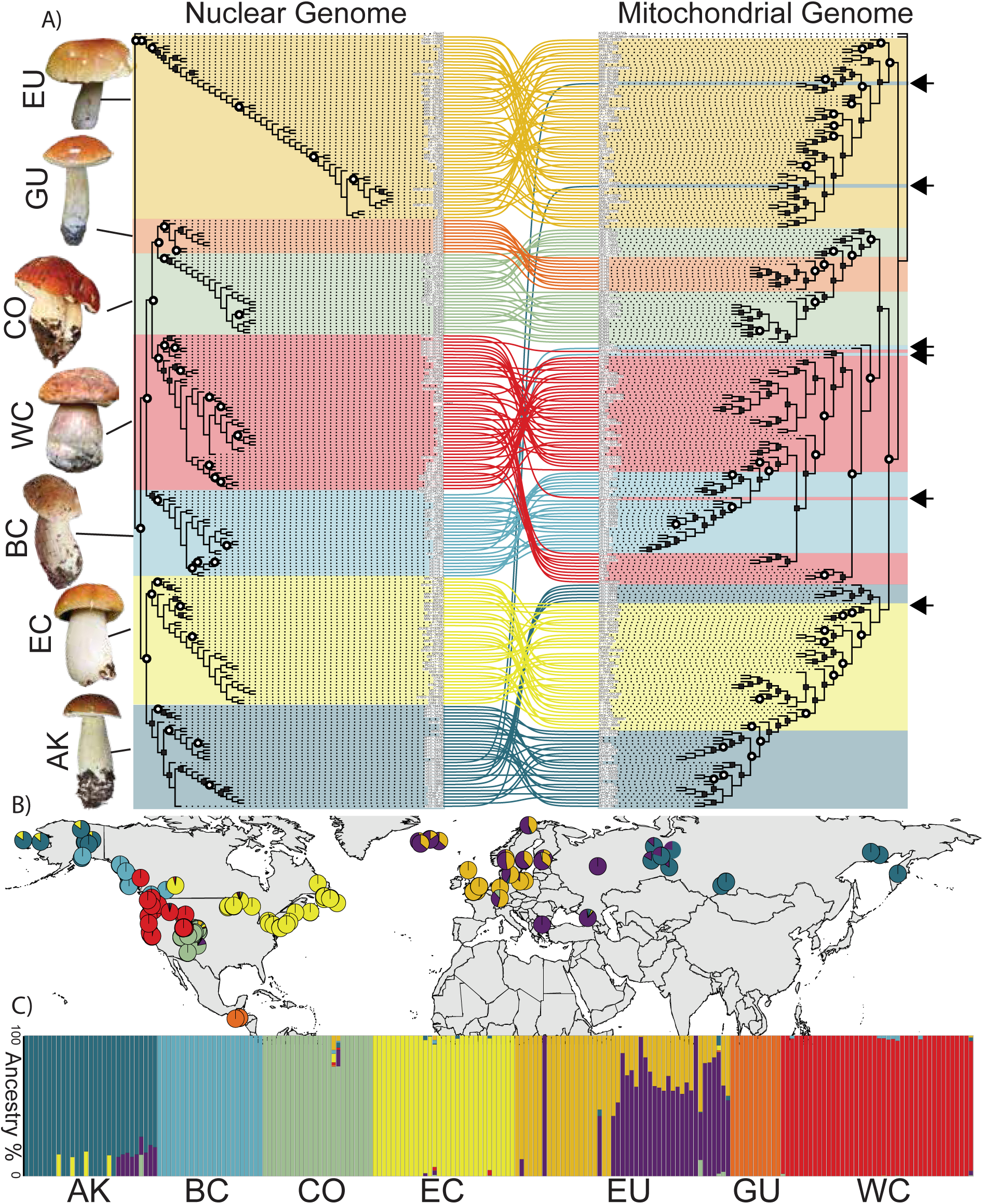
Cyto-nuclear discordance and admixture analysis of *B. edulis*. A) Cophylogenetic analysis of nuclear genome species tree (left) and mitochondrial genome tree (right). Cophylogeny was produced with the “co.phylo” comand of the R package “phytools”. White circles on phylogeny indicate nodes with 100% nodal support (quartet support on left, bootstrap support on right), and stars indicate nodes with at least 80% nodal support. Pictures left of phylogeny are examples of the phenotypic diversity found within *B. edulis*. From bottom to top YSU-5404 (AK), BD294 (EC), UT567 (WC), BD942 (CO), KST39 (GU), KST15 (BC), All photos were taken by B. Dentinger, except YSU-5404 (used with permission from Ni_2_n_8_a Filippova) and KST15 (from K. Tremble). It is important to note that phenotypes do not cleanly segregate into lineages. B) Distribution of estimated admixture across the global distribution of *B. edulis*. C) Barchart of admixture proportions per sample per lineage.

### 2.2 Analysis of genome-wide admixture and divergence, suggest strong reproductive barriers among BC-WC lineages

To determine whether contemporary hybridization among lineages can lead to introgression, we genotyped single nucleotide polymorphisms (SNPs), and estimated levels of genome wide admixture and divergence (Fst). In addition, we implemented two empirical tests utilizing variations of f4 statistics to validate whether signatures of admixture were the product of introgression or incomplete lineage sorting. Admixture analysis revealed evidence of population structure within the EU lineage, and generally low levels of admixture shared among lineages except within the AK lineage (Fig. 1)C, Fig. S1). When conducting admixture analyses with K>6 we found that the EU lineage was split into two populations (Fig. S1), a high-latitude Fennoscandia and Icelandic population, and a mainland Europe and United Kingdom population, confirming other recent reports [31]. Interestingly, this European population structure mirrors that found in other basidiomycetes [10]. Within these two populations of the EU lineage numerous individuals exhibit substantial admixture from both population clusters indicating gene-flow and the lack of strong speciation barriers. Despite parapatric and near sympatric distributions we found little to no evidence for shared admixture among lineages, except within the AK lineage (Fig. 1 B,C), confirming earlier results [13]. Four individuals of the AK lineage exhibit admixture from the EC lineage, and nine exhibit admixture from the EU lineage, specifically from the high-latitude population. Importantly, admixed individuals within the AK lineage were not distributed randomly. Alaskan individuals of the AK lineage show evidence of admixture with the EC lineage, western Russian individuals of the AK lineage show evidence of admixture with the EU lineage, and central/eastern Russian individuals exhibit no admixture from either. If the presence of admixture in AK was solely due to the retention of ancestral polymorphisms, we would expect ancestral alleles to be randomly distributed in descendant populations and not concentrated geographically near lineage boundaries. Altogether, admixture analyses suggests that genomic introgression is present among the AK-EU-EC lineages, while the BC-WC lineages exhibit little, despite signatures of ongoing hybridization.

Among lineages, the EC and AK lineages showed the least differentiation (AK-EC Fst=0.146, Table S6, Fig. S2), and the AK-EC comparison produced by far the lowest values of Fst for both lineages, corroborating evidence for ongoing hybridization and introgression. In contrast, the GU lineage exhibited the highest differentiation with every other lineage (Mean Fst = 0.265), with the exception of CO (Fst=0.191). For both the BC and WC lineages, the BC-WC comparison exhibited the least differentiation (Fst=0.180) potentially corroborating evidence for abundant historic introgression. However this level of differentiation was only slightly lower than other comparisons (EC-WC = 0.186, BC-EC = 0.188), which may again indicate that introgression among BC and WC is limited today.

To empirically test for introgression rather than incomplete lineage sorting we utilized both the qpAdm algorithm from admixtools2 and Patterson’s D (ABBA-BABA) statistic from Dsuite. qpAdm is an implementation of the f4 statistic and models a target population as a mixture of several proxy ancestry sources, and has been shown to consistently identify true signatures of admixture among complex demographic histories [32, 33]. Specifically we tested whether the AK lineage could be modeled as being an admixed population of the EC and EU lineages, or rather some other combination of lineages. The only model that could not be rejected was the model where EC and EU contribute to admixture in AK (Table S7), with EC contributing 80.8% percent of admixture. D statistics also support ongoing introgression rather than incomplete lineage sorting (Table S8). The EC-AK-EU trio exhibited the highest D-statistic and f4 ratio of all comparisons with an ABBA/BABA ratio significantly supporting introgression (D=0.174, Z=3.46, P=0.00053, f4=0.343 **??**). The BC-WC-EC trio was also highly significant (D=0.143, Z=2.6655, P=0.0077, f4=0.164), corroborating patterns of cyto-nuclear discordance among these lineages.

### 2.3 Whole genome mapping of 23 reference genomes reveals low conservation of synteny and abundant structural variation that is rarely shared

To determine whether large lineage-specific genomic structural variants contribute to post-zygotic reproductive barriers, preventing introgression among lineages, we used whole genome mapping of the 22 reference quality genomes generated here against the publicly available European chromosomal reference genome [31] to identify structural variants larger than 1Kbp. We predicted 3,147 duplications (5.9 Kbp average length), 2,442 translocations (13.0 Kbp average length), and 252 predicted inversions (25.0 Kbp average length) (Table S9, Table S10, Fig. S3). Surprisingly few inversions or translocations were shared between individuals of the same lineage: 322 duplications (10% of total), 11 inversions (4% of total), and 124 translocations (5.1% of total). Moreover, no inversion and only one small translocation (1753bp, in BC lineage) was found in all three genomes of a single lineage. Fig. 2A highlights the striking lack of conservation of predicted structural variants among lineages of *B. edulis*; only three structural variants within chromosome nine are found in more than one genome. To complement this analysis, we used orthologous gene mapping between our new near-chromosomal reference genomes from the AK and WC lineage and the European Chromosomal reference (Fig. 2B.) to identify structural variants. With this approach we identified larger and more abundant translocations and inversions with the genome from the AK lineage than the WC lineage, including three translocations greater than 500Kbp and one greater than 1Mbp.

**Figure 2:**
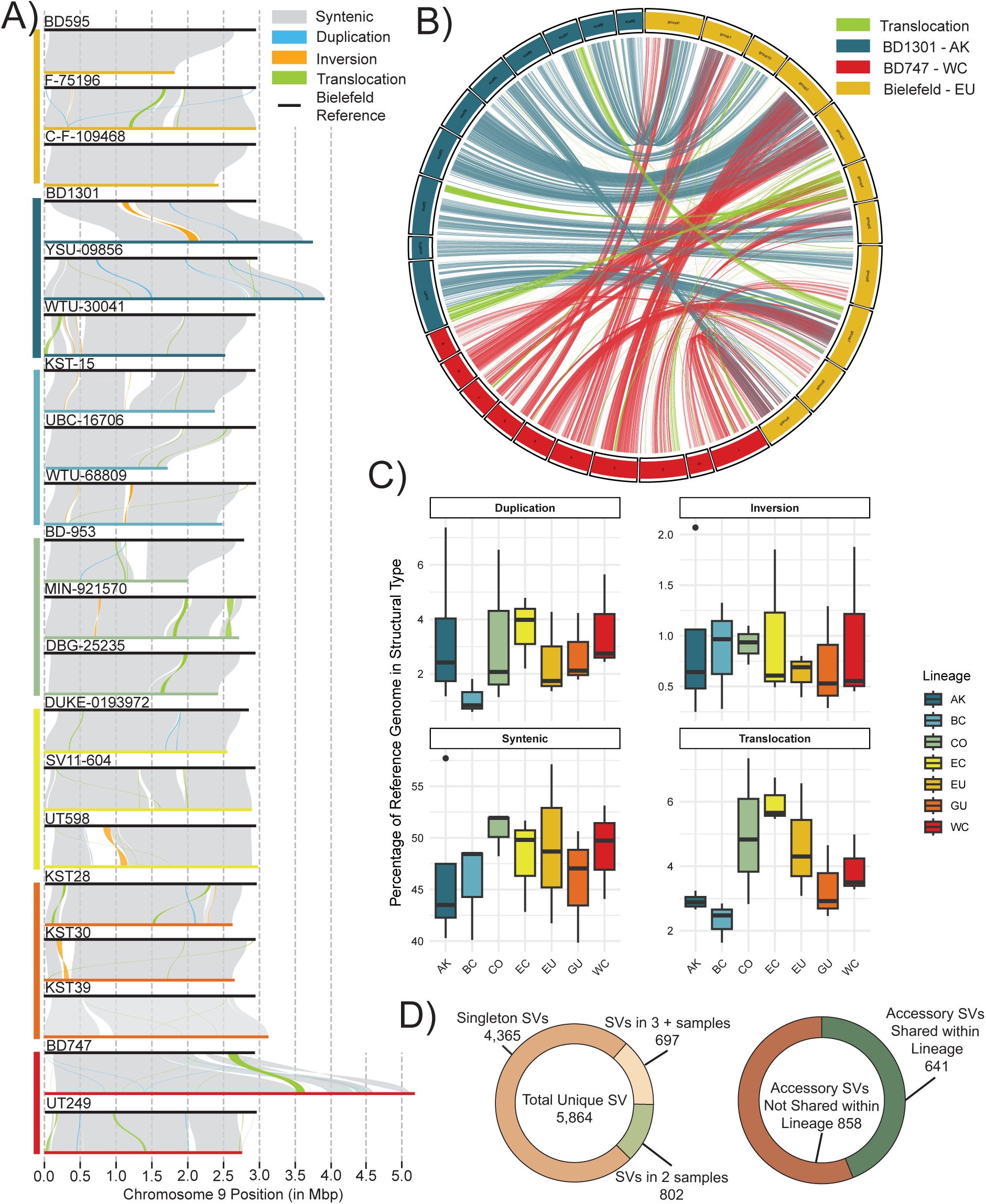
Identification of large genomic structural variation and synteny among lineages of B. edulis. A) Result of whole genome mapping of 22 reference genomes against chromosome 9 of the *B. edulis* European Chromosomal Reference genome, which highlights the ample structural variation present in B. edulis.. Genomes are grouped by lineage with color bars on left corresponding to lineages from Fig. 1 and Fig. 2 C. Only three of all structural variants within chromosome 9 are found within more than one genome, and no structural variant is found in all three genomes of a lineage. B) Identification of large structural variants our two deeply sequenced genome assemblies against the *B. edulis* European Chromosomal Reference using orthologous gene mapping rather than whole genome mapping. Tracks in green indicate likely translocations. Translocations are more numerous and substantially larger between the BD1301 from the AK lineage than BD747 from the WC lineage.

To estimate the extent to which genomic synteny is conserved among and within lineages of *B. edulis*, we performed whole genome mapping against the European chromosomal reference genome using the SYRI pipeline and orthologous gene mapping between pairs of genomes to identify blocks of syntenic regions. We found strikingly low levels of synteny across both analyses. On average, only 47% of the European chromosomal reference genome was found to be syntenic with each other genome after whole genome mapping (Fig. 2 C). Estimation of pairwise synteny using orthologous gene blocks found even lower levels of synteny; 37.5% of each genome on average (Fig. S4). Surprisingly, similar levels of synteny were found among samples of the same lineage as between samples of different lineages. For example, members of the EU lineage exhibit a lower average degree of synteny with the European chromosomal reference than either the CO, EC, or WC lineages, despite being members of the same lineage as the reference. This indicates that on average, genomic synteny is under weak conservation within *B. edulis*.

### 2.4 The *B. edulis* super-pan-genome reveals unprecedented genomic diversity

To corroborate the above analyses of genomic structural variation and synteny, we constructed the “super pan-genome” of *B. edulis* and determined whether lineages with signatures of both hybridization and introgression (AK-EU-EC) exhibit greater pan-genomic similarity than lineages without introgression (BC-WC and CO-GU). Pan-genomic approaches use *de novo* genome assemblies to identify the gene families that are “core”— families that are present in all genomes, and that are likely necessary for key molecular and ecological functions of the species — and “accessory” – gene families that are present in some but not all genomes.

Across 200 samples we identified a total of 30,719 orthologous gene clusters (Fig. 3B, Table S11, Table S12). Of these orthologous groups, 2,120 (6.9% of total) were singletons (found exclusively in one genome), 10,399 (33.9%) were rare (found in at least two, but less than five genomes), 12,597 (41%) were accessory (found in at least five genomes, but less than 95% of all genomes), and 7,256 (23.6%) were core (found in at least 95% of genomes). Surprisingly, only four orthologous gene groups were lineage-specific and present in at least of 90% samples within that lineage (one group in BC lineage and three in GU, see supplemental results for details). To determine whether lineages with signatures of hybridization/introgression have similar genomic compositions, we clustered all specimens according to their accessory protein profiles using PCoA (presence/absence of orthologous gene group only, gene-copy number agnostic). We identified four clear clusters that correlate with geography and not phylogeny or introgression (Fig 3D). The BC-WC, and GU-CO lineages formed two nearly indistinguishable pan-genomic clusters clearly separated from the other lineages. By contrast, the EC lineage was segregated into a single isolated cluster, and the AK-EU lineages formed distinct but slightly overlapping pan-genomic clusters.

**Figure 3:**
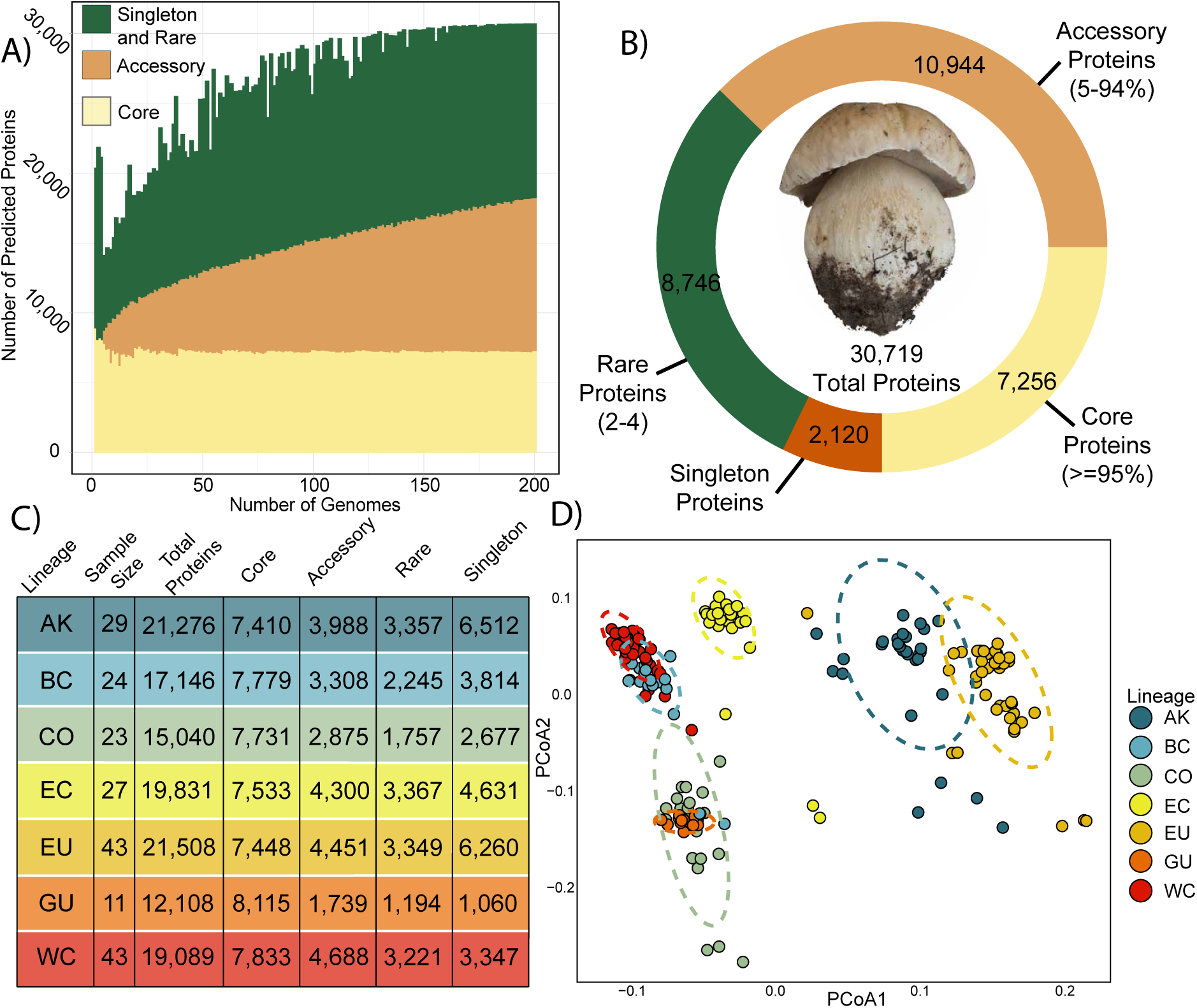
The global super pan-genome of *B. edulis*. A) The abundance of pan-genome enzyme categories as a function of number of genomes in the analysis. B) The abundance of core (>=95% presence), accessory (N=5-94% presence), rare (N=2-N=4) and singleton (N=1) within the *B. edulis* pan-genome. C) The abundance of core and accessory proteins found within each individual lineage of *B. edulis*. D) Principal coordinate analysis (PCoA) of 22 *B. edulis* genomes based on accessory protein profiles. Elipses represent 95% confidence intervals.

### 2.5 Identifying whether lineages of *B. edulis* differ in ecologically relevant gene families

To determine whether lineages with signatures of contemporary hybridization and/or introgression are more ecologically similar we characterized the abundance and composition of three groups of ecologically important gene families: 1) secreted and non-secreted carbohydrate active enzymes (CAZYmes) and peptidases (MEROPS) used for biodegredation and substrate modification (termed biodegredation repertoire), 2) small secreted proteins (SSPs) and membrane-bound nutrient transporters which are widely implicated in host associations (termed host-association repertoire), and 3) specific gene families involved in abiotic stress (e.g. osmotic stress, starvation stress).

Within the biodegredation repretoire, we identified 262 CAZymes on average (112 secreted CAZymes on average) in *B. edulis* genomes, which is substantially fewer than has been found within other Boletales EMF taxa [34, 35] (. 4A, Table S13-S15). We also identified further reductions in enzymes targeting cellulose/hemicellulose, such as AA3 oxidoreductases (average = 9.6), and GH5 hemicellulases (average = 14.0). However, *B. edulis* appears to possess a greater number of peptidases on average (mean = 503) compared to other EMF taxa [36]. Within *B. edulis*, we found significant differences in the abundance of both CAZymes and peptidases among lineages. The GU lineage showed consistent contractions in most degradation enzyme families and possessed the fewest CAZymes and peptidases in total (P < 0.01, F=4.311, DF=6, one-way ANOVA, Fig. S5, Fig. S6). To determine whether the lineages differ in their overall biodegredation profiles we used PCoA to cluster specimens based on their composition and abundance of CAZymes and peptidases (Fig. 4B). We found that the AK-EU-EC lineages comprise a single overlapping cluster while all other lineages exhibit little to no overlap with any other lineage.

**Figure 4:**
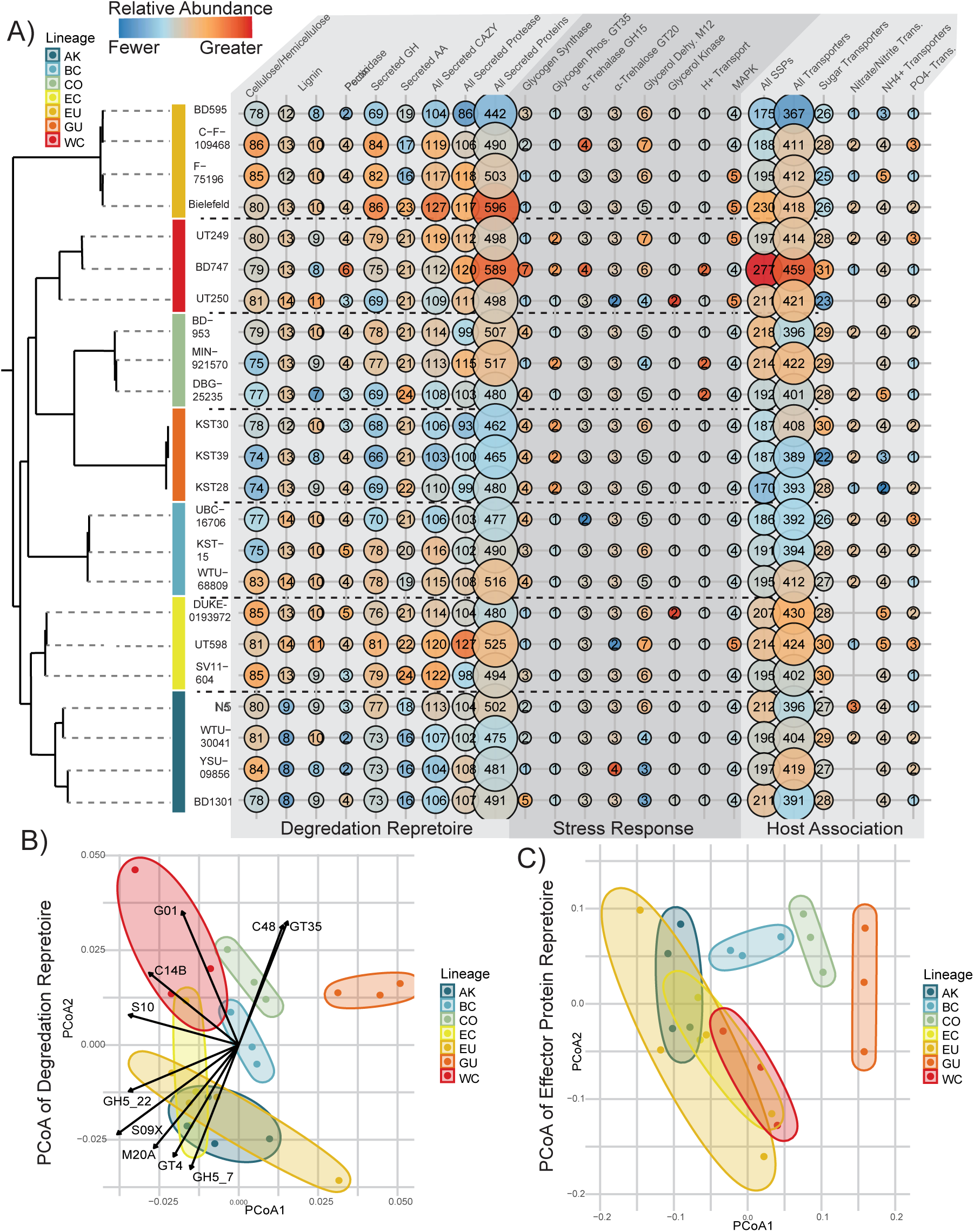
Analysis of ecologically relevant gene families in *B. edulis*. A) Abundance of biodegredation, stress-response, and host-association gene repertoires across the seven lineages of *B. edulis*. The size of the bubbles indicates the number of proteins within each protein family. The lack of a bubble indicates the complete loss of an enzyme family. The color of the bubble indicates the relative abundance within *B. edulis*, where blue indicates relative contractions and red indicates relative expansions. Names below chart indicate either primary substrate of enzyme class (e.g. chitin) or name of enzyme if substrate not well characterized in EMF (e.g glucoamylase). B) Results of the principal coordinate analysis (PCoA) of biodegredation repertoires. Elipses do not indicate and confidence intervals. Arrows indicate variables significantly contributing variables. C) Results of the PCoA of Small Secreted Protein profiles. Elipses do not indicate and confidence intervals.

Within the host-association repertoire, we identified 393 unique orthologous SSP groups and 202 SSPs on average within a single genome with no significant differences in abundance being present among the lineages (Table S16-S17). Of these SSP groups only 29 were found in all 23 reference genomes, and only one SSP in the CO lineage was lineage specific and found within all three genomes. To determine whether the lineages differ in their SSP profiles we used PCoA to cluster specimens (Fig. 4C). We found substantial SSP overlap among the AK-EU-EC and WC lineages while the other lineages exhibit no overlap with any other lineage. For membrane-bound transporters we found 407 proteins on average distributed across 88 protein families, and while no differences among lineages were significant, the EC and WC lineages tended to possess a greater abundance (Fig. 4A, Table S18-S19). Within the most well characterized classes of transporters involved in host association such as sugar transporters, nitrate/nitrite transporters, ammonium transporters, and phosphate transporters [37, 38], we found no evidence of lineage-specific differences or clear patterns of similarity among specific lineages. Lastly, we found limited evidence of signatures of differentiation in the gene copy number of genes involved in abiotic stress responses. Copy numbers were either largely conserved, or widely variable with no clear patterns of shared expansion or contraction.

## 3 Discussion

### 3.1 Porcini consist of seven globally distributed lineages with contrasting patterns of hybridization and introgression

Identifying contemporary hybridization in natural populations is a difficult task, even when the organisms in question are readily accessible and display identifiable hybrid phenotypes. Soil-borne fungi are often neither. Thus, we relied on phylogenomic approaches to identify signatures of hybridization. Phylogenetic discordance between nuclear and mitochondiral genomes can arise after hybridization events [39] and has been widely investigated across most eukaryotic life to identify instances of hybridization between long-diverged taxa [40, 41, 42, 43], and even in diverse basidiomycete fungi [44, 45, 46]. Mitochondrial inheritance is predominately uniparental in fungi [47]. When biparental inheritance is present, it is effectively uniparental as different mitochondria are spatially segregated in the mycelium of an individual [43, 48]. Thus, patterns of cyto-nuclear discordance in Fungi can reflect differences in mitochondiral and nuclear genotypes. Our results suggest that the CO-GU, AK-EC, AK-EU, and BC-WC lineages have experienced historical introgression leading to substantial cyto-nuclear discordance. This is supported by significant D statistics within trio comparisons containing these lineages that would indicate the presence of introgression (Table S8). Similar patterns of discordance between nuclear and mitochondrial genomes has been identified in both ascomycete [49, 50] and basidiomycete [51, 44] fungi, which suggests that this approach could be a broadly applicable and powerful tool to identify patterns of historic introgression. However, only the AK-EU, AK-EC, and BC-WC lineages exhibited individuals with differing nuclear and mitochondrial genotypes, which is indicative of contemporary hybridization as has been shown in Heterobasidion spp. [52, 53]. This suggests that recent reproductive barriers have arisen between the CO and GU lineages, or that sampling in potential hybrid zones is insufficient to capture rare hybridization events. It is worth noting that we identified mitochondrial hybrids between the AK-EU-EC lineages despite limited sampling within the AK-EU hybrid zone, and no sampling within the AK-EC hybrid zone (Fig. 1B). This suggests that hybridization among these lineages is frequent enough that signatures of hybridization, or actual hybridization, can extend beyond regions of high contact.

Hybridization among the diverged phylogenetic species of *B. edulis* appears to be somewhat common, yet hybridization events do not necessarily lead to introgression. Numerous genomic and ecological forces can prevent potential hybrids from ever producing viable offspring. Analysis of genome-wide admixture and differentiation from the AK-EU-EC lineages, indicates ongoing introgression, despite an estimated 2.1 million years of divergence [13]. These lineages exhibit the highest degree of admixture, lowest pairwise population divergence (Fst), and qpAdm and D statistics support introgression rather than solely incomplete lineage sorting. In contrast, we found little to no indication that hybridization between the BC and WC lineages is leading to introgression. Despite substantial sampling within the geographic hybrid zone, we identified no individuals with substantial admixture between the BC and WC lineages, and no large reduction of population divergence (Fst) as found in the AK and EC lineages. This suggests that stronger reproductive barriers are present among the BC and WC lineages than the AK-EC-EU lineages. It is important to note we found no evidence of lineage-specific divergence or loss of synteny within the primary mating locus of *B. edulis* (MATa) (Supp. Results, Fig. S7, Fig. S8, Table S21) that could contribute these reproductive barriers. Altogether, our results imply that within *B. edulis*, there are loose controls on the capacity to hybridize, but rather, strong controls on whether hybridization leads to introgression. Some other notable basidiomycete fungi also exhibit this pattern, such as *Cryptococcus spp.* which readily hybridize (both *in vitro* and in natural populations) yet often produce sterile hybrids [54, 44, 55, 56]. However, other basidiomycetes can display the opposite pattern, such as *Trichaptum abietinum* where *in vitro* hybrids cannot be formed from sympatric isolates, demonstrating strong pre-zygotic boundaries [10]. Just as in other eukaryotes, the formation of reproductive barriers in fungi is not a deterministic process, and different barriers will manifest depending on the demographic and ecological context [1].

### 3.2 Exceptional structural variation persists within and among lineages of Porcini

Genomic architecture plays an indisputably important role in evolution, and large structural changes are widely implicated in the formation of reproductive barriers [57, 20, 58]. Here, we found that *B. edulis* possesses substantial large structural variants, both within and among lineages, but these are rarely shared between different individuals and are largely found within a single genome. Surprisingly, we found that only one small translocation is found within all three specimens of a lineage (BC). We also found no relationship between the degree of synteny and the presence/absence of hybridization. The AK and EU lineages exhibit signatures of both hybridization and introgression, yet the AK lineage exhibits by far the lowest average degree of synteny with the European chromosomal reference. For example, YSU-09856, an individual from the AK lineage with a relatively high degree of genomic admixture from the EU lineage, exhibits the third lowest degree of synteny among all samples (40.2%, Fig. 2C). These results suggest that natural populations of fungi harbor extensive structural variation, and that this variation appears to play a limited role in limiting introgression among species.

We found that *B. edulis* possess one of the most diverse super pan-genomes among eukaryotes with the lowest proportion of core proteins so far identified [59, 60]. Even within individual lineages, six out of seven *B. edulis* lineages contain a lower percentage of core proteins (mean = 44.5%, Fig. 3C) than has been previously identified in other fungi [61, 62]. *B. edulis* is one of the more widely distributed eukaryotic taxa on earth, inhabiting a vast array of habitats. Even within a single lineage, specimens of *B. edulis* can be found at sea-level in Californian coastal pine forests, and at over 3000 meters in alpine spruce and fir forests (Tremble personal observations). We hypothesize that the vast ecological amplitude exhibited by *B. edulis* is associated with its exceptional pan-genomic diversity. Despite this diversity, we found substantial clustering among lineages in pan-genomic profile; clusters that corresponded with geography and not phylogeny or presence/absence of introgression. For example, the BC and WC lineages exhibit total pan-genomic overlap, indicating highly similar genome composition. While these two lineages do hybridize, indicated by the presence of cyto-nuclear discordance and three individuals with hybrid mitochondrial genotypes, they exhibit limited to no signatures of introgression. This suggests that strong postzygotic boundaries exist between BC and WC in the absence of genome-wide structural variation. Further, the CO and GU lineages also exhibit total pan-genome overlap, yet show even less evidence of introgression than the BC and WC lineages. In contrast, the lineages with strong signatures of both hybridization and introgression, AK-EU/AK-EC, exhibit limited to no pan-genomic overlap. This corroborates our finding that AK-EU exhibits the lowest degree of synteny among all lineages (Fig. 2C). Thus structural variation is at best a weak barrier for introgression among long diverged lineages in *B. edulis*.

### 3.3 *B. edulis* as a group may specialize on chitin degradation, yet lineages still exhibit ecological differentiation

Though EMF play a dynamic role in soil nutrient cycling [63, 64, 65], it remains unclear whether their biodegradation repertoires reflect specific ecologies or preferred habitats [66]. Here, we found that *B. edulis* exhibits a substantially reduced biodegradation repertoire compared to most other EMF regardless of geography, including further losses of cellulolytic enzymes. Despite these overall losses we found a generally high abundance of GH18 (glycoside-hydrolase family 18) chitinases. Dead fungal biomass, or “necromass”, represents a substantial pool of carbon and nitrogen in soils [67], yet can be a difficult target for decomposition due to the often abundant presence of highly aromatic melanin [68, 69]. Despite this recalcitrance, it has been consistently shown that EMF are abundant on fungal necromass, and can rapidly incorporate carbon from this pool [70]. Not only does *B. edulis* contain abundant secreted GH18 chitinases, but we also find an expansion of AA6 (Auxiliary-Activity family 6) 1,4-benzoquinone reductases (Table S13), which are intra-cellular enzymes involved in the biodegradation of aromatic compounds [71, 72], and which may provide a mechanism to facilitate melanin decomposition. While this capacity needs to be empirically validated, our results are consistent with a shift in the ecology of *B. edulis* away from the “mining” of nutrients from plant derived sources towards the utilization of fungal necromass.

Despite this shared pattern, we still found evidence of ecological differentiation among lineages of *B. edulis*, namely within the biodegredation gene repertoire. The most clear example is the tropical GU lineage, which exhibits additional contractions in most biodegredation gene families (Fig. 4A), and an altogether distinct profile (Fig. 4B). It has long been proposed that tropical soils represent a unique habitat for EMF compared to temperate soils [73]. This is because high decomposition rates lead to rapid nutrient cycling and low levels of soil organic matter in tropical soils [74]. Moreover, a greater abundance of sunlight hours and precipitation in tropical latitudes increases the photosynthetic capacity of EMF hosts, increasing the abundance of readily available carbon that can be allocated to EMF [75, 76]. These factors may reduce the need for EMF to enzymatically liberate nitrogen and other nutrients from soil organic matter, or the need to acquire carbon from non-host sources during periods of low sunlight, subsequently yielding a contraction in the decomposition machinery of the GU lineage. While more cryptic than the GU example, other lineages also exhibit signatures of ecological differentiation. The WC lineage exhibits a significant expansion of metallopeptidases, which have been shown to be important enzymes for soil-organic matter degradation by EMF [77, 78], and a greater abundance of G1 peptidases (Fig. 4B) which are known to be active at acidic pH [79]. As a group, the AK-EU-EC lineages were differentiated from the rest of *B. edulis* by significant differences in several biodegredation families including - mannanases (GH5_7) (Fig. 4B). While both plants and fungi contain mannan polysacharrides [80], the expanded -mannanases in the AU-EU-EC lineages contain predicted secretion peptides (Table S15) and we believe are likely targeting extracellular pant mannan.

We also found that the BC and WC lineages differ in their host-association gene repertoire, suggested by no overlap in SSP clustering (Fig. 4C). While single SSPs have been shown to be decisive for EMF colonization [81], it is unclear whether the difference in host-association gene repertoires among BC and WC represents a shift in preferred host, or simply novel mechanisms for interacting with shared hosts. For example, a recent study of the ectomycorrhizal genus *Suillus* found little difference in total abundance of SSPs between host specialists and host generalists [82]. Host associations of EMF are difficult to empirically determine and no clear differences in preferred hosts exist among *B. edulis*. It is thought that *B. edulis* from North America are primarily restricted to gymnosperms, with the vast majority of observations recorded in Pinaceae-dominated forests and with confirmed angiosperm associations being relatively rare (MyCoPortal. 2024. http://www.mycoportal.org/portal/index.php. Accessed on October 3). However, we have one WC specimen in our dataset that was collected in a mono-dominant stand of *Populus trichocarpa* (DUKE-351166, Table S1) which indicates that the capacity to colonize diverse hosts exists within the WC lineage, albeit rarely. Thus, we hypothesize that these differences in host-association relevant gene families among the BC and WC lineages likely does not represent a clear shift in preferred host. Rather, we hypothesize that these differences represent alternative mechanisms for interacting with shared hosts.

### 3.4 Pattern of hybridization and introgression correlate with ecological similarity and not structural variation

Hybridization is common across all domains of eukaryotic life, but appears to be particularly common in fungi. However, some fungi display a substantially greater tendency for hybridization than others and it is unclear what intrinsic characteristics facilitate or prevent hybridization. Why do some fungal taxa hybridize while others do not? We found evidence that hybridization among diverged species within the *B. edulis* species complex is somewhat common, particularly among the AK-EU, AK-EC, BC-WC lineage pairs. These lineages exhibited multiple “mitochondrial-hybrids” indicative of recent hybridization. However, only the AK-EU and AK-EC lineage pairs exhibit signatures of ongoing introgression, such as the presence of admixture, low population divergence (Fst), and significant qpadm and D statistics. To understand why hybridization leads to introgression in the AK-EU-EC lineages, but not within the BC-WC lineages, we identified whether genomic structural variation could be contributing to reproductive boundaries within the BC-WC lineages. Yet, we found that the BC and WC lineages comprise a single pan-genomic cluster and no evidence of large structural variants fixed within either lineage. By contrast the AK lineage exhibits the lowest degree of synteny with our EU reference genome, and the AK-EU-EC lineages exhibit the lowest degree of pan-genomic similarity of all lineages. Thus, our results indicate that at the scale of large natural populations the presence or absence of structural variation alone cannot predict whether two long diverged lineages will exhibit signatures of introgression.

Instead, we find that patterns of introgression correlate with ecological similarity. The AK-EU-EC lineages possess the most similar profiles of ecologically relevant gene families, including shared expansions of extracellular mannanases and overlapping SSP profiles. Whereas the BC and WC lineages, and in particular the CO and GU lineages, exhibit clear differences in both host association and biodegredation profiles. While these ecologically relevant gene families and their role in organic-matter degradation and host association has been well characterized in other highly diverse fungi [83, 84, 85], their role in the specific ecology of *B. edulis* will need to be validated in subsequent work. However, we believe that our results indicate the presence of ecological differentiation among the BC-WC and CO-GU lineages; or at least, a degree of differentiation that was not indicated by analyses of genomic structural variation. Moreover, *B. edulis* is not the only organism, let alone fungus, where ecology appears to be the primary mechanism mediating introgression. Within the *Heterobasidion annosum* species complex, a group of mushroom-forming plant pathogens, taxa that exhibit similar host preference readily hybridize and are highly interfertile despite 30-41 million years of divergence [86, 87, 88]. By contrast, *Heterobasidion* taxa that exhibit clear differences in host preference rarely hybridize; and when rare hybrids are found, they are found on a shared host [89]. Just as in the analogous *Heterobasidion*, when we identify signatures of introgression among the species of *B. edulis*, it is among the taxa that likely exhibit the most ecological similarity, and not the taxa that possess the most similar genome structure. Altogether, these results highlight the role that ecology plays in reinforcing species boundaries in fungi. Moreover, we believe our results suggest that some mechanisms which clearly cause reproductive barriers *in vitro*, function with far less clarity in natural populations of fungi.

## 4 Methods

### Specimen Collection, Sequencing, and Assembly

253 *Boletus edulis* specimens were collected from a global distribution using museum collections, augmented with targeted field sampling campaigns in Alaska, Utah, Germany, the United Kingdom, and Guatemala (Table S1). We also included recently generated telomere-telomere reference genome of *B. edulis* [31]. Genomic DNA was extracted from dried dried hymenophore tissue using an optimized version of Monarch® Genomic DNA Purification Kit (New England Biolabs). Samples were sequenced using a combination of paired-end sequencing on the Illumina MiSeq, HiSeq, and Novaseq sequencing platforms. Short-read genome assemblies were construced using four different assembly algorithms, (see supplemental methods for full list) and the assembly with the highest recovery of BUSCO genes in the “basidiomycota_odb10” dataset was retained as the final assembly for each sample. Lastly, we identified changes in predicted genome size and genome-wide heterozygosty using the kmer-based estimation approach implemented in Genomescope with Jellyfish used for kmer identification.

### Confirmation as “Boletus edulis” and Lineage Placement

To confirm the specimens as *B. edulis* and to classify them into lineages, we used a summary coalescent phylogenomic approach. From each genome, we identified 1764 highly-conserved single copy orthologs using BUSCO [90] with the “basidiomycota odb 10” dataset. Sequence alignment of retained orthologs was performed using MAFFT v7.397 [91], and maximum-likelihood gene-trees were inferred using IQ-TREE v2.0.3 [92]. A summary coalescent species tree was constructed from the resulting gene trees using the astral-hybrid algorithm from from ASTER (v1.16) according to author recommendations [93].

### Reference Quality Whole Genome Sequencing and Genome assembly

After lineage determination we re-sequenced and generated reference quality genomic assemblies for 22 individuals, representing at least three individuals per lineage. 20 of 22 individuals were re-sequenced at moderate coverage (2°0-50x) using the Oxford Nanopore Flongle flowcell (VR10.1) and hybrid genome assemblies were generated with MaSuRCA (V4.0.9) [94]. The remaining two specimens we re-sequenced at high coverage (250x) using the Oxford Nanopore Minion (R10) to generate pseudo-chromosomal assemblies. Assemblies were generated for each specimens using most publicly-available single-molecule genome assemblers (see supplemental methods). A final consensus assembly was generated using the RagTag pipeline [95]. After all 22 reference genome assemblies were created, duplicate sequences were removed using the purge_dups pipeline (V1.2.3 https://doi.org/10.1093/bioinformatics/btaa025), and assemblies were polished using three successive rounds of PILON (V1.24)[96].

### Gene Prediction within all genome assemblies

For short-read genome assemblies with a BUSCO recovery score of at least 90% and our 22 reference quality genome assemblies we conducted gene prediction. First, contigs less than 500bp in length and those taxonomically classified as “Bacteria” with MMseqs2 were purged from each assembly. Next, repetitive content we predicted using the EDTA [97] wrapper script, and repatative content was “softmasked”. Gene prediction and initial functional annotation was performed using the Funannotate (v1.8.9) gene prediction wrapper script according to proscribed practices [98].

### Cophylogenetic Analysis of the Nuclear and Mitochondiral Genome to Assess Cyto-Nuclear Discordance

To construct the mitochondrial phylogeny, we aligned filtered and adapter trimmed sequencing reads of all *B. edulis* individuals to the mitochondrial genome of *B. edulis* accessed from Mycocosm (Boled5 genome, [99]) with Bowtie2 (v2.2.6) [100]. Variants were called using BCFtools, and a consensus sequence was generated using the mpileup command. A maximum-likelihood tree based on the unpartitioned alignment was inferred with IQ-TREE v2.0.3. Cophylogenetic comparison between mitochondrial tree and nuclear species tree was performed using the “cophylo” function of the R package Phytools (1.5-1).

### Estimating Admixture, Divergence, and ABBA-BABA Among Lineages of *B. edulis*

Single nucleotide polymorphisms (SNPs) were identified with the GATK 3 pipeline using Illumina short reads aligned to the EU chromosomal reference genome “BolEdBiel_h2” [31]. SNPs missing in more than 20% of samples, with a mean depth of coverage less than 10 and more than 100, and with an allele present in less than 3 samples were removed from the dataset with VCFtools (v0.1.15) [101]. Lastly, samples with more than 75% missing data were removed from the dataset. Pairwise genome wide Weir and Cockerham’s Fst was calculated using VCFtools command –weir-fst-pop. To identify signatures of genomic admixture we first conducted linkage disequilibrium (LD) filtering using plink2 (https://www.cog-genomics.org/plink/2.0/credits) across 5kb windows, with 1kb window jump, and a max r^2^ value of 0.2. Admixture coefficients were calculated with the program ADMIXTURE [102] using values of K from five to nine. To validate whether signatures of admixture were the product of introgression and not solely incomplete lineage sorting we used two separate empirical tests. Specific models of admixture were empirically tested using the qpAdm software implemented in admixtools2 (v2.0.4) [103]. qpAdm is an implementation of the f4 statistic and models a target population as a mixture of several proxy ancestry sources, and has been shown to consistently identify true signatures of admixture among complex demographic histories [32, 33]. Second, we conducted Patterson’s D (ABBA-BABA) tests to identify signatures of introgression across all possible trios using Suillus luteus (DOB657 Resequencing, SRR22259488) as the outgroup with the Dsuite package [104].

### Identifying genomic synteny

The 22 Reference genomes generated here were aligned to the EU chromosomal reference genome “BolEdBiel_h2” using minimap2 (V2.28 [105]) with the option “-ax asm5”. From these alignments we identified structural variants greater than 1Kbp using the SYRI package (V1.7.0 [106]). Structural variants of the same predicted type with beginning and end positions within 1000bp of each other were classified as “shared” variants. We also identified blocks of syntenic orthologous genes, sets of orthologous genes with the same order forward and reference, between pairs of genomes using the “super-pangenome” constructed below. Using these syntenic orthologous gene blocks we first visualized the “gene links” between our two best reference genome assemblies (BD747 and BD1301) and our chromosomal reference genome (Bielfeld) using a custom script (see code availability) and the “circlize” R package [107].

### Construction of the *B. edulis* “Super-Pangenome“

Orthologous genes were identified with Orthofinder using the “–diamond-ultra-sens” option [108] from predicted proteins from 200 short-read genome assemblies and 23 reference genomes. Genes were classified as “core” (present in 95% of genomes), “accessory” (found in at least five genomes, but less than 95% of all genomes), “rare” (found in at least two genomes, but less than five), and “singleton”. To visualize lineage clustering based on accessory gene profile, we converted all accessory proteins to presence/absence counts (gene copy number agnostic), and implemented principal coordinates analysis (PCoA, cmdscale function of stats R package) using a Bray-Curtis distance matrix (calculated with Vegan R package).

### Degradation and carbohydrate interactive enzymes

Carbohydrate activate enzymes (“CAZymes”), were identified from our 23 reference genomes using the run_dbCan4 program [109]. Only proteins receiving at least 2 significant hits were retained. In addition we also identified intracellular and extracellular peptidases using HMMSCAN (V3.3.2) searches of protein sequences against the MEROPS peptidase database [110]. Significant hits were classified as Evalue < 1e^−15^, and only the hit with the lowest Evalue was retained. We used PCoA to cluster specimens by their total CAZyme and peptidase repertoire, utilizing Bray curtis distances calculated by the “vegdist” function of the vegan R package (v2.6.4). To identify specific enzyme families contributing to the segregation of individual samples we calculated correlations between variables and PCoA axes using the “envfit” command of the vegan R package (v2.6.4) with 999 permutations.

### Identifying unique host interactions

Small secreted proteins (SSPs) potentially involved in host association were predicted from our 23 *B. edulis* reference genomes using two AI large-language model algorithms 1) tmbed (V1.0 [111], and 2) deeploc (V2.1 [112]). Proteins were classified as “secreted” if they contained a predicted secretion signal peptide from both algorithms, predicted to lack a transmembrane helix from the tmbed algorithm, and predicted to be “extracellular soluable” from the deeploc algorithm. Only secreted proteins smaller than 300 amino acids were retained. To visualize differences in SSP composition and gene copy number (SSP repertoire) we performed PCoA as above. Transporters were identified by using a HMMSCAN search against the reference HMMs from the Transporter Classification Database (TCDB [113]). Significant hits were classified as Evalue < 1e^−15^, and only the hit lowest Evalue was retained.

### Stress Response Pathways

We identified abundance of genes within, 1) starvation stress response, 2) osmotic stress response, 3) general conserved stress response pathways. Specifically we searched for, 1) the glycogen synthesis (IPR008631) and metabolism (CAZY GT35 = glycogen-phosphorylase) pathways, which are implicated in starvation and energy limitation, 2) glycerol production (Aerobic glycerol-3-phosphate dehydrogenase = MEROPS M12,), glycerol regulation (HOG1 = IPR038783), glycerol transportation, and aquaporin pathways which are all intricately involved in the osmotic stress response, 3) and the generic mitogen activate kinase pathways (MAPK, IPR003527).

## 7 Acknowledgments

The authors would like to acknowledge the many institutions and individuals who provided specimens for use in this work (See Table S1 for a full list of collaborating holding institutions). We would like also like to thank Tiffany Do and Sariah VanderVeur for their effort in extracting genomes for this work.

## 6 Data Availability

All short-read genome sequences, are publicly available on the Short Read Archive and Genbank under the BioProect accession no. PRJNA1010140 and BioSample ID numbers for each collection used in this study are found in Table S1. Genome assemblies and predicted protein sequences for all short-read only genome sequences are deposited on Dryad and will be accessible by request or once published (https://doi.org/10.5061/dryad.hmgqnk9r4). Assemblies, predicted protein sequences, and genome annotations for the 22 references are deposited on NCBI GENOME portal under the BioProject accession no. PRJNA1216376. Custom scripts used for analyses are deposited on github (https://github.com/KeatPorcini/Porcini-pan-genomics) and are freely accessible by request or once published, see supplementary methods for the name of each script used within for each analysis.

