## Supplementary material for "From populations to pan-genomes: investigating the role of ecology and genomic architecture in maintaining species boundaries in the porcini mushroom, *Boletus edulis*": Fig. S: Tremble_et_al_SupplementaryMethodsResultsFigures_25feb25.pdf

### 1 Supplementary Expanded Methods

#### 1.1 Sample Collection

253 *Boletus edulis* specimens were collected from a global distribution using museum collections, augmented with targeted field sampling campaigns in Alaska, Utah, Germany, the United Kingdom, and Guatemala (Table S1). Specimen vouchers of new collections are deposited in the mycological collection of the Garrett Herbarium at the University of Utah. We also included recently generated telomere-telomere reference genome of *B. edulis* [29]. In addition, we included publicly-available genome data from *Paxillus involutus* (Batsch) Fr., *Paxillus adelphus* J.-P. Chaumeton, Gryta, Jargeat P.-A. Moreau, *Hydnomerulius pinastri* (Fr.) Jarosch & Besl [26] retrieved from the JGI Mycocosm Portal [17], and the taxon sister to *B. edulis*, *Boletus reticuloceps* (M. Zang, M.S. Yuan M.Q. Gong) Q.B. Wang Y.J. Yao (NCBI accession PRJNA702108 [58, 11]) [57] as outgroups.

#### 1.2 Short-Read Whole Genome Sequencing and Genome Assembly

Genomic DNA was extracted from dried specimens using 10mg of hymenophore tissue. Tissue was first physically ground to a fine powder using 1 x 3.0 mm and 10-15 x 0.5 mm stanless steel beads in a 2.0 mL screwcap tube by shaking in a BeadBug$^{TM}$ homogenizer set at 400 rmp for 30 seconds. Fully homogenized tissue was then used for DNA purification using the "Protocol for Extraction and Purification of Genomic DNA from Tissues" protocol for extraction and purification of genomic DNA from tissues in the Monarch® Genomic DNA Purification Kit (New England Biolabs T3010). DNA extract quality was assessed for quality (260:230 nm and 260:280 nm wavelengths) using a NanoDrop 1000 spetrophotometer (Thermo Scientific) and DNA integrity was assessed using agarose gel electrophoresis. Samples were sequenced using a combination of paired-end sequencing on the Illumina MiSeq, HiSeq, and Novaseq sequencing platforms at the Genomics Core, University of Utah (Table S2).

#### 1.3 Specimen Collection, Sequencing, and Assembly

253 *Boletus edulis* specimens were collected from a global distribution using museum collections, augmented with targeted field sampling campaigns in Alaska, Utah, Germany, the United Kingdom, and Guatemala (Table S1). We also included recently generated telomere-telomere reference genome of *B. edulis* [29]. Genomic DNA was extracted from dried dried hymenophore tissue using an optimized version of Monarch® Genomic DNA Purification Kit (New England Biolabs). Samples were sequenced using a combination of paired-end sequencing on the Illumina MiSeq, HiSeq, and Novaseq sequencing platforms. Short-read genome assemblies were construced using four different assembly algorithms, (see supplemental methods for full list) and the assembly with the highest recovery of BUSCO genes in the "basidiomycota_odb10" dataset was retained as the final assembly for each sample. Lastly, we identified changes in predicted genome size and genome-wide heterozygosty using the kmer-based estimation approach implemented in Genomescope with Jellyfish used for kmer identification.

Raw sequencing reads from 253 specimens were quality-filtered and adapter-trimmed using fastP v0.20.1 [7] with default settings. Short-read genome assemblies were produced from quality-filtered reads using four methods, 1) SPAdes v3.15.0 [4] in "–isolate" mode, 2) SPAdes in "–meta" mode, 3) MEGAHIT in generic assembly mode ("–min-count 3 –no-mercy"), and 4) MEGAHIT in "–sensitive" mode. COMPLEASM, a faster and more accurate re-implementation of BUSCO, was used to assess assembly quality of all four genome assemblies, and the assembly with the highest recovery of BUSCO genes in the "basidiomycota_odb10" dataset was retained as the final assembly for each sample. Lastly, we identified changes in predicted genome size and genome-wide heterozygosty using the kmer-based estimation approach implemented in Genomescope with Jellyfish used for kmer identification.

#### 1.4 Confirmation as "Boletus edulis" and Lineage Placement

To confirm the specimens as *B. edulis* and to classify them into lineages, we used a summary coalescent phylogenomic approach and partitioned Maximum likelihood analysis to estimate branch lengths.Across our dataset, we identified 1,757 highly-conserved single copy orthologs BUSCO [59] genes derived from the "basidiomycota odb 10" lineage dataset. Orthogroups were filtered using SEGUL 0.22.1(Handika Esselstyn, 2024)to only include genes found in at least 75% of samples, leading to a final count of 1,610 BUSCO genes used for downstream analysis. Sequence alignment of filtered orthologs was performed using MAFFT v7.397 [24] with the L-INS-i algorithm, and maximum-likelihood gene-trees were inferred using IQ-TREE v2.3.6 [39] with automatic model selection in ModelFinder [23] and ultrafast bootstrapping (BS, [21]) with 1000 replicates. A summary coalescent species tree was constructed from the resulting gene trees using Weighted ASTRAL (wASTRAL) algorithm (Zhang  Mirarab, 2022) from the Accurate Species Tree EstimatoR (ASTER) v1.16 package (https://github.com/chaoszhang/ASTER/blob/master/tutorial/wastral.md). Branch lengths for summary coalescent species tree were estimated using IQ-TREE v2.3.6 [39] with a concatenated and partitioned data set of filtered orthologous produced using SEGUL 0.22.1(Handika  Esselstyn, 2024).

#### 1.5 Reference Quality Whole Genome Sequencing and Genome assembly

After lineage determination we sought to generate reference quality genomic assemblies for 22 individuals (23 including the telomere-telomere reference previously generated [29]), representing at least three individuals per lineage. Genomic DNA was extracted using a modified phenol/chloroform isolation method (see [55]) optimized for high molecular weight DNA. 20 of 22 individuals were re-sequenced at moderate coverage ($\tilde{2}$0-50x) using the Oxford Nanopore Flongle flowcell (VR10.4.1) and and hybrid genome assemblies were generated with MaSuRCA (V4.0.9) [63] using both Illumina short-reads and Nanopore long-reads. This approach was chosen for the majority of the samples intended for reference-genome generation as it is the most cost-effective and high-throughput way to produce numerous highly contiguous genome assemblies. For the remaining two specimens we sought to produce near-chromosomal level genome assemblies for fine comparisons of synteny with the newly generated chromosomal reference genome. To this end, we re-sequenced these two specimens at exceptionally high coverage ( 250x) using the Oxford Nanopore Minion (R10). Individual assemblies were generated for each specimens using most publicly-available single-molecule genome assemblers, including Necat [8], Nextdenovo [42], Smartdenovo [33], FLYE [27], Canu [28], and MaSuRCA [63] with a minimum read-length of 1kb. A final consensus assembly was generated using the RagTag pipeline [2] by retaining scaffold patterns produced by at least two separate assemblers. After all genome assemblies, duplicate sequences were removed using the purge_dups pipeline (V1.2.3 https://doi.org/10.1093/bioinformatics/btaa025), and assemblies were polished using three successive rounds of PILON (V1.24)[56].

#### 1.6 Gene Prediction within all genome assemblies

For subsequent analyses we conducted gene prediction within all high-quality short-read genome assemblies with a BUSCO recovery score of at least 90% and also our 22 reference quality genome assemblies. First, we removed potential contaminant contigs in our genome assemblies by taxonomically classifying contigs with the MMseqs2 "easy-taxonomy" function and the full NCBI "NR" database. All contigs assigned to "Bacteria", including descendants, were removed form the assembly as well as contigs below 500 base-pairs in length. Next, we predicted repetitive content within each assembly using the EDTA [44] prediction wrapper script (with "–anno 1" options), and used these predictions to create a "soft masked" genome assembly using the "maskfasta" command of Bedtools (https://bedtools.readthedocs.io/en/latest/). Gene prediction was performed utilizing the Funannotate (v1.8.9) gene prediction wrapper script according to proscribed practices [45]. To help guide gene prediction, we used our previously published training set parameters constructed from *B. edulis*, (BD747, [54]), as well as previously generated mRNA sequences ([54]) mapped to each genome with hisat2 (V2.2.1) [25]. General functional annotation of predicted proteins from the 22 reference-quality genomes was conducted using Interproscan (V5.69 https://doi.org/10.1093/bioinformatics/btu031) and Eggnog-Mapper V(2.1.12 10.1093/nar/gky1085) and the Funannotate annotate wrapper script.

##### 1.7 Cophylogenetic Analysis of the Nuclear and Mitochondiral Genome to Assess Cyto-Nuclear Discordance

To construct the mitochondrial phylogeny, we aligned filtered and adapter trimmed sequencing reads of all *B. edulis* individuals to the mitochondrial genome of *B. edulis* accessed from Mycocosm (Boled5 genome, [40]) with Bowtie2 (v2.2.6) [30]. Variants were called using BCFtools [10], and a consensus sequence was generated using the mpileup command. A maximum-likelihood tree based on the unpartitioned alignment was inferred with IQ-TREE v2.0.3. Cophylogenetic comparison between mitochondrial tree and nuclear species tree was performed using the "cophylo" function of the R package Phytools (1.5-1). To verify whether patterns of nuclear mitochondrial discordance are the product of incomplete lineage sorting or introgression we used the HyDE program [6], which has explicitly been developed to test for patterns of hybridization in phylogenomic data, on our aligned Mitochondrial sequences.

##### 1.8 Cophylogenetic and Synteny Analysis of Mating-type Loci in *B. edulis*

To further verify that signatures of introgression are contemporary and not simply remainders of historic gene flow we determined whether mating loci exhibit any signatures of lineage specific divergence that would indicate sexual incompatibility. To this end we 1) identified the genes within the MATa locus within our 23 reference quality genomes and characterized patterns of synteny, and 2) compared phylogenies of the homeo-domain loci (HD1 and HD2) and the nuclear genome to identify whether lineage specific divergence exists.

To determine whether structural and genetic variation within primary mating locus of *B. edulis* (MATa), may be contributing to reproductive barriers we identified the composition and synteny of the homeodomain MATa locus within our 22 reference genomes plus the telomere-telomere reference. First we used the HD1 and HD2 loci from the European telomere-telomere reference [29] as queries in a blastp search against predicted proteins within each other reference genome with an evalue cutoff of 1e-15. We then recorded the orientation and predicted annotation of (using gff3 files from each genome) of the two genes upstream of HD1 and all genes downstream of HD2 until GLG2 (if present). Homology of other predicted genes with the MATa locus with the European reference was assessed using blastp with an evalue cutoff of 1e-15.

To investigate if sexual incompatibility could be acting as a driver of lineage segregation in *B. edulis*, we performed a cophylogenetic analysis between the homeodomain from the fungal mating-type locus HD1 and the species tree of *B. edulis*. Fungi exhibit non-associative mating, where sexual compatibility is determined by genetic diversity in a set of mating-type genes (MAT loci). Sexual reproduction in Fungi occurs randomly in a conspecific population when mating-type loci are sufficiently different. If lineages are segregated through sexual incompatibility, then lineage relationships of the mating-type loci should mirror that of the species tree. Exonerate version 2.4.0 1 was used to extract homologs of the mating-type locus HD1. Homologs were extracted from assembled scaffolds using the "–protein2genome" and "–bestn 2" flags to account for the dikaryotic nature of the samples. Comparison of unique and identical alleles within and between populations was performed using SeqKit (v2.7.0) with the flags "–common –by-seq –ignore-case" [51] Query sequences of HD1 from *Rhizopogon salubrious* coding sequences (CDS) (ON315867.1) were used as they were the most closely related publicly available sequences. Each allele of the HD1 homologs was aligned using the multiple sequence alignment program MAFFT (version 7.475) with the parameters "–maxiterate 1000 –localpair –reorder". Multiple sequence alignments were used as input for gene tree construction using IQTREE, as above. Cophylogenetic comparison between the mating-type gene trees and *B. edulis* species tree was performed using the "cophylo" function of the R package Phytools (1.5-1)

#### 1.9 Estimating Admixture, Divergence, and ABBA-BABA Among Lineages of *B. edulis*

To genotype genome-wide single nucleotide polymorphisms (SNPs) within *B. edulis*, we first aligned adapter-trimmed and quality-filtered short reads to the the EU chromosomal reference genome "BolEdBiel_h2" [29] with Bowtie2 (v2.2.6) [30]. We then identified SNPS from these alignments using the GATK 4 pipeline according to author instructions [3]. SNPs missing in more than 20% of samples, with a mean depth of coverage less than 10 and more than 100, and with an allele present in less than 3 samples were removed from the dataset with VCFtools (v0.1.15) [9]. Lastly, samples with more than 30% missing data were removed from the dataset. To prevent the inclusion of putative clones in all analyses we used the –relatedness2 function of VCFtools to identify specimens with a high probability of clonality. If two or more sequences were identified as clonal, using the threshold of 0.354 according to author recommendations, we only retained the sequence with the best assembly BUSCO score, and other sequences were removed from the dataset for all downstream analyses.

Pairwise genome wide Weir and Cockerham's Fst was calculated using VCFtools command –weir-fst-pop. To identify signatures of genomic admixture we first conducted linkage disequilibrium (LD) filtering using plink2 (https://www.cog-genomics.org/plink/2.0/credits) with the command "–indep-pairwise" across 5kb windows, with 1kb window jump, and a max $r^2$ value of 0.2. LD pruning resulted in a final total of 52,738 SNPs. Admixture coeficiants were calculated with the program ADMIXTURE [1] using values of K from five to nine. Admixture coefficients were plotted in R using ggplot2. To validate whether signatures of admixture were the product of introgression and not solely incomplete lineage sorting we used two separate empirical tests. Specific models of admixture were empirically tested using the qpAdm software implemented in admixtools2 (v2.0.4) [37]. qpAdm is an implementation of the f4 statistic and models a target population as a mixture of several proxy ancestry sources, and has been shown to consistently identify true signatures of admixture among complex demographic histories [20, 60]. Second, we conducted Patterson's D (ABBA-BABA) tests to identify signatures of introgression across all possible trios using Suillus luteus (DOB657 Resequencing , SRR22259488) as the outgroup with the Dsuite package [38].

#### 1.10 Identifying Genomic Synteny

To directly quantify the degree of synteny that exists between any two lineages we performed 1) whole genome mapping of our reference-quality genomes against our chromosomal reference and predicted structural variants from the alignments and 2) identified syntenic gene blocks between genomes using the gene orthology prediction from the super-pangenome analyses above.

To characterize synteny from whole genome alignments we first used minimap2 (V2.28 [31]) with the option "-ax asm5" to align genomic scaffolds from each reference quality assembly to our Bielefeld chromosomal reference genome. From these alignments we re-scaffolded our assemblies into "pseudo-chromosomes" using the "chroder" command of the SYRI package (V1.7.0 [15]). It is important to note that by scaffolding our assemblies against a single reference we are likely underestimating structural variation present within *B. edulis*, yet scaffolding is necessary for prediction of large structural variants via whole genome mapping. After scaffolding, we used the SYRI package to identify large structural variants (minimum 1000 base pairs) within each genome against the Bielefeld chromosomal reference. Alignments were visualized with the PLOTSR package (V1.1.0 [16]). Lastly, we sought to identify whether predicted structural variants were found only within a single genome or were broadly distributed across *B. edulis*. To this end we used a custom script to identify structural variants whose start and end position within the Bielefeld reference genome were within 1000 base pairs of another structural variant of the same type within another genome ("Boletus_edulis_parse_SYRI.R", https://github.com/KeatPorcini/Porcini-pan-genomics).

To characterize synteny by identifying blocks of syntenic genes we identified orthologous genes within all reference genomes using the "super-pangenome" constructed below. Using these orthologous genes we first visualized the "gene links" between our two best reference genome assemblies (BD747 and BD1301) and European refernece genome using a custom script ("plot_circos_BD747_BD1301_EUref https://github.com/KeatPorcini/Porcini-pan-genomics) and the "circlize" R package [18]. To compare the size and quantity of syntenic blocks between all reference genomes we identified blocks of genes that were predicted as orthologous between two genomes and exhibited the same order (forward or reverse) in both genomes ("syntenic_blocks_from_orthologs.R", https://github.com/KeatPorcini/Porcini-pan-genomics).

#### 1.11 Construction of the *B. edulis* "Super-Pangenome"

Orthologous genes were identified with Orthofinder using the "–diamond-ultra-sens" option [12] from predicted proteins from 200 short-read genome assemblies with BUSCO recovery >= 90% and 23 reference genomes. All genes not placed in orthogroups (excluding singleton genes placed in their own orthogroup) were removed from subsequent analyses. Genes were classified as "core" (present in 95% of genomes), "accessory" (found in at least five genomes, but less than 95% of all genomes), "rare" (found in at least two genomes, but less than five), and "singleton" using a custom script "Boletus_edulis_pangenomics_19sep2024.R", https://github.com/KeatPorcini/Porcini-pan-genomics). To visualize lineage clustering based on accessory gene profile, we converted all accessory proteins to presence/absence counts (gene copy number agnostic), and implemented principal coordinates analysis (PCoA, cmdscale function of stats R package) using a Bray-Curtis distance matrix (calculated with Vegan R package).

#### 1.12 Identification of of the "Biodegredation" Repretoire of *B. edulis*

Carbohydrate activate enzymes ("CAZymes"), were identified from our 23 reference genomes using the run_dbCan4 program [62]. Only proteins receiving at least 2 significant hits were retained. After CAZyme and peptidase prediction we sought to identify specific differences in ecological tendencies among lineages of *B. edulis* by grouping gene families by predicted carbohydrate substrate (specifically Cellulose/hemicellulose, ligin, pectin, and chitin, see Table S14 for exact groupings), and using ANOVA to test for differences in gene copy number. In addition we also identified intracellular and extracellular peptidases using HMMSCAN (V3.3.2) searches of protein sequences against the MEROPS peptidase database [48]. Significant hits were classified as Evalue $< 1e^{-15}$, and only the hit with the lowest Evalue was retained. We used PCoA to cluster specimens by their total CAZyme and peptidase repertoire, utilizing Bray curtis distances calculated by the "vegdist" function of the vegan R package (v2.6.4). To identify specific enzyme families contributing to the segregation of individual samples we calculated correlations between variables and PCoA axes using the "envfit" command of the vegan R package (v2.6.4) with 999 permutations.

##### 1.13 Identifying unique host interactions

To identify whether lineages of *B. edulis* differ in their likely capacity to colonize and interact with varied hosts, we characterized the abundance and diversity of small secreted proteins that may facilitate host-microbe communication, and proteins involved in sugar, nitrogen, and phosphorus transport that likely facilitate the ectomycorrhizal partnership between fungus and host. Secreted proteins were predicted from the proteomes of our 23 *B. edulis* reference genomes using two AI large-language model algorithms 1) tmbed (V1.0 [5], and 2) deeploc (V2.1 [43]). Proteins were classified as "secreted" if they were predicted to contain a secretion signal peptide from both algorithms, predicted to lack a transmembrane helix from the tmbed algorithm, and predicted to be "extracellular soluable" from the deeploc algorithm. Small secreted proteins (SSPs) were characterized as secreted proteins smaller than 300 amino acids. To identify whether SSPs where sample or lineage specific we used the "super-pangenome" constructed above to identify orthologous gene clusters containing predicted SSPs. To visualize differences in SSP composition and gene copy number among lineages we performed PCoA as above of all specimens using their "SSP repretoire". Transporters were identified by using a HMMSCAN search against the reference HMMs from the Transporter Classification Database (TCDB [49]). Significant hits were classified as Evalue $< 1e^{-15}$, and only the significant hit with the lowest Evalue was chosen for each predicted protein.

##### 1.14 Stress Response Pathways

Lastly we sought to identify potential differences in ecological tendencies among lineages of *B. edulis* by characterizing the abundance of genes within well characterized fungal stress responses. To this end, we sought to identified abundance of genes within, 1) starvation stress response, 2) osmotic stress response, 3) general conserved stress response pathways. Specifically we searched for, 1) the glycogen synthesis (IPR008631) and metabolism (CAZY GT35 = glycogen-phosphorylase) pathways, which are implicated in starvation and energy limitation [41], 2) glycerol production (Aerobic glycerol-3-phosphate dehydrogenase = MEROPS M12, ), glycerol regulation (HOG1 = IPR038783), glycerol transportation, and aquaporin pathways which are all intricately involved in the osmotic stress response, 3) and the generic mitogen activate kinase pathways (MAPK).

### 2 Supplementary Expanded Results

#### 2.1 Short-Read Whole Genome Sequencing and Genome Assembly

Whole genome sequencing of 253 collections of *B. edulis* resulted in 10,091,414 Illumina read pairs on average yielding an average genome coverage of 75.7x ( 40Mbp average genome size). The genome assembler Megahit in generic assembly mode ("–min-count 3 –no-mercy") was the best assembler overall (median BUSCO recovery = 96.37, average = 88.98), then Megahit in ultra-sensitive present mode (median BUSCO recovery = 96.26, average = 88.93), then metaSpades (median BUSCO recovery = 95.63, average = 89.04), and finally Spades in "isolate" mode (median BUSCO recovery = 49.86, average = 55). Of all four assemblies for each collection, the best assembly was produced by Megahit in generic mode 115 times, Megahit sensitive 73 times, Spades in "isolate" mode 39 times, and metaSpades 20 times. Of all the best assemblies for each collection median BUSCO score was 96.6, average BUSCO score was 90.8.

GenomeScope analysis of genome size and genome heterozygosity identified significant differences among lineages (Fig. ??C,D). Average genome size across all of *B. edulis* was 40.0 Mbp, and average genome heterozygosity was 1.29%. The GU lineage was the least heterozygous (0.623%) and had the largest predicted genome size (47.0 Mbp) ($P < 0.05$, one-way ANOVA, Fig. ??C)

#### 2.2 Reference Genome Sequencing and Assembly

Hybrid-sequencing produced 20 highly-contiguous reference genomes (mean N50 = 344 Kbp, mean number of scaffolds = 570, Table S3). In addition, we sequenced two samples at deep coverage with Oxford Nanopore long reads, BD747 of the WC lineage and BD1301 of the AK lineage, and produced two additional pseudo-chromosomal reference assemblies (BD747 N50 = 2.2Mbp, BD1301 N50 = 2.5Mbp). Average assembly length across all assemblies was 39.58 Mbp (median = 39.9 Mbp). Repetitive content predicted with the EDTA pipeline varied from a minimum of 13.67% (UBC-16706, 4.72 Mbp repetitive content) to a maximum of 31.03% (BD1301, 14.00 Mbp repetitive content), with an average of 23.07% (9.36 Mbp repetitive content). The average number of predicted proteins across all reference genomes (including the publicly available European reference genome) was 11,802.

#### 2.3 Confirmation as "Boletus edulis" and Lineage Placement

Summary coalescent analysis of 1,610 gene-trees with ASTER revealed seven reciprocally monophyletic *B. edulis* clades with 100% statistical node support that share a most recent common ancestor with the Asian endemic *B. reticuloceps* (Fig. 1A,). Six of these lineages directly correspond to the six lineages previously identified [54], and hereafter will be referred to as AK (Alaska/Siberia, BC (British Columbia), CO (Colorado), EC (Eastcoast North America), EU (Europe), and WC (Westcoast North America). The seventh lineage is a novel group that is exclusively comprised of tropical *B. edulis* specimens from Guatemala and will henceforth be referred to as GU (Guatemala).

#### 2.4 Cophylogenetic Analysis of the Nuclear and Mitochondiral Genome to Assess Cyto-Nuclear Discordance

Cophylogenetic analysis of the nuclear genome species tree and mitochondrial genome tree reveled substantial patterns of cyto-nuclear discordance (Fig. 1A.) Three lineages that appear monophyletic in the nuclear species tree (AK, WC, and CO) were polyphyletic in the mitochondrial tree, with well-supported nodes (BS>0.75) leading to separated clusters. Analysis of introgression within the mitochondrial genome using HyDE suggests the discordance among the nuclear and mitochondrial genomes are not solely the product of incomplete lineage sorting. Hyde detected significant signatures of introgression (P<0.05) in 15 trio comparisons, and all lineages polyphyletic in the mitochondrial tree were included within at least one trio pair with significant signatures of introgression (Table S4). For example, the BC lineage was placed within the WC lineage in the mitochondrial tree, and HyDE identified a trio comparison containing EC, BC, WC as possessing highly significant signatures of introgression (Z=2.1, P=0.017). In addition, we identified at least five individuals with hybrid genotypes; in these cases, an individual was placed within the monophyletic lineage consistent with its collection location in the nuclear species tree, but was placed within another lineage in the mitochondrial tree with significant nodal support. Two individuals from the AK lineage were placed in the EU mitochondrial group, two individuals from BC lineage in the WC lineage, and one from the WC lineage was placed within the BC lineage. Interestingly, one individual from the EC lineage, was placed outside of *B. edulis* and sister to *B. reticuloceps* in the mitochondrial tree. We excluded this individual from all subsequent analyses.

#### 2.5 Analysis of genome-wide admixture, divergence in *B. edulis*, and f4 statistics to verify patterns of introgression

Admixture analysis revealed evidence of population structure within the EU lineage, and generally low levels of admixture shared among lineages except within the AK lineage (Fig. 1C, Fig. S1). At values of K 5-6, the CO and GU, and BC and WC lineage pairs were each clustered into a single admixture group, while all other lineages were placed into their own admixture groups. As the AK and EC lineage pair exhibits substantially lower pairwise population divergence (Fst, see below) than CO-GU or BC-WC we would have expected AK-EC to cluster at lower values of K rather than CO-GU or BC-WC. This surprising clustering likely reflects the bias produced by using a single reference genome from the European lineage to identify SNPs, leading to decreased mapping rates of divergent sequences from other lineages. Yet, when the analysis was conducted with K=8, we found that all lineages were placed into their own admixture group. When conducting admixture analyses with K>6 we found that the EU lineage was split into two populations (Fig. S1), a high-latitude Fennoscandia and Icelandic population, and a mainland Europe and United Kingdom population, confirming other recent reports [29]. Interestingly, this European population structure mirrors that found in other basidiomycetes [36]. Within these two populations of the EU lineage numerous individuals exhibit substantial admixture from both population clusters indicating geneflow and the lack of strong speciation barriers. Despite parapatric and near sympatric distributions we found little to no evidence for shared admixture among lineages, except within the AK lineage (Fig. 1 B,C), confirming earlier results [54]. Four individuals of the AK lineage exhibit admixture from the EC lineage, and nine exhibit admixture from the EU lineage, specifically from the high-latitude population. Importantly, admixed individuals within the AK lineage were not distributed randomly. Alaskan individuals of the AK lineage show evidence of admixture with the EC lineage, western Russian individuals of the AK lineage show evidence of admixture with the EU lineage, and central/eastern Russian individuals exhibit no admixture from either. If the presence of admixture in AK was solely due to the retention of ancestral polymorphisms, we would expect ancestral alleles to be randomly distributed in descendant populations and not concentrated geographically near lineage boundaries. Altogether, admixture analyses suggests that genomic introgression is present among the AK-EU-EC lineages, while the BC-WC lineages exhibit little, despite signatures of ongoing hybridization.

Among lineages, the EC and AK lineages showed the least differentiation (AK-EC Fst=0.146, Table S6, Fig. S2), and the AK-EC comparison produced by far the lowest values of Fst for both lineages, corroborating evidence for ongoing hybridization and introgression. In contrast, the GU lineage exhibited the highest differentiation with every other lineage (Mean Fst = 0.265), with the exception of CO (Fst=0.191). For both the BC and WC lineages, the BC-WC comparison exhibited the least differentiation (Fst=0.180) potentially corroborating evidence for abundant historic introgression. However, this level of differentiation was only slightly lower than other comparisons (EC-WC = 0.186, BC-EC = 0.188), which may again indicate that introgression among BC and WC is limited today.

To empirically test for introgression rather than incomplete lineage sorting we utilized both the qpAdm algorithm from admixtools2 and Patterson's D (ABBA-BABA) statistic from Dsuite. qpAdm is an implementation of the f4 statistic and models a target population as a mixture of several proxy ancestry sources, and has been shown to consistently identify true signatures of admixture among complex demographic histories [20, 60]. Specifically we tested whether the AK lineage could be modeled as being an admixed population of the EC and EU lineages, or rather some other combination of lineages. The only model that could not be rejected was the model where EC and EU contribute to admixture in AK (Table S7), with EC contributing 80.8% percent of admixture. D statistics also support ongoing introgression rather than incomplete lineage sorting (Table S8). The EC-AK-EU trio exhibited the highest D-statistic and f4 ratio of all comparisons with an ABBA/BABA ratio significantly supporting introgression (D=0.174, Z=3.46, P=0.00053, f4=0.343 **??**). The BC-WC-EC trio was also highly significant (D=0.143, Z=2.6655, P=0.0077, f4=0.164), corroborating patterns of cyto-nuclear discordance among these lineages.

#### 2.6 Characterizing the abundance of distribution of large structural variants

To determine whether large lineage-specific genomic structural variants contribute to post-zygotic reproductive barriers, preventing introgression among lineages, we used whole genome mapping of the 22 reference quality genomes generated here against the publicly available European chromosomal reference genome [29] to identify structural variants larger than 1Kbp. We predicted 3,147 duplications (5.9 Kbp average length), 2,442 translocations (13.0 Kbp average length), and 252 predicted inversions (25.0 Kbp average length) (Table S9, Table S10, Table S22). Surprisingly few inversions or translocations were shared between individuals of the same lineage: 322 duplications (10% of total), 11 inversions (4% of total), and 124 translocations (5.1% of total). Moreover, no inversion and only one small translocation (1753bp, in BC lineage) was found in all three genomes of a single lineage. Variants were more or less equally distributed among chromosomes when controlling for chromosome size (Inversion = 6.8/Mbp, Translocation = 60.9/Mbp, Duplications = 78.62/Mbp, Table S22, Fig. S5), yet average variant size varied widely among chromosomes. For example, chromosome 1 exhibited a median inversion size of 20,225 Bp, almost six times larger than the median inversion size on chromosome 6 (3,476 Bp).

#### 2.7 Identifying translocations within three near chromosomal reference genomes using orthologous gene mapping

To identify patterns of synteny among chromosomal level reference genomes, we used orthologous gene mapping between our two new near-chromosomal reference genomes from the AK (BD1301) and WC (BD747) lineages and the European Chromosomal reference (Fig. 2B.) to identify structural variants. When compared with the European reference genome we identified numerous translocations within both BD747 and BD1301. BD1301 exhibited fewer but substantially larger translocations than BD747, three translocations greater than 500Kbp and one greater than 1Mbp (1.82Mbp, Translocation from chromosome 7 to chromosome 4). Moreover, the entirety of chromosome 4 from the European reference appears to have been split and fused with scaffold 1 and scaffold 2 of BD1301. In contrast BD747 possessed more numerous, but smaller translocations with the exception of one 654.8 Kbp translocation from chromosome 5 to chromosome 3 (Table S23).

#### 2.8 Identifying conservation of synteny within all reference genomes of *B. edulis*

To estimate the extent to which genomic synteny is conserved among and within lineages of *B. edulis*, we performed whole genome mapping against the European chromosomal reference genome using the SYRI pipeline and orthologous gene mapping between pairs of genomes to identify blocks of syntenic regions. We found strikingly low levels of synteny across both analyses. On average, only 47% of the European chromosomal reference genome was found to be syntenic with each other genome after whole genome mapping (Fig. 2C). Estimation of pairwise synteny using orthologous gene blocks found even lower levels of synteny; 37.5% of each genome on average (Fig. S4). Surprisingly, similar levels of synteny were found among samples of the same lineage as between samples of different lineages. For example, members of the EU lineage exhibit a lower average degree of synteny with the European chromosomal reference than either the CO, EC, or WC lineages, despite being members of the same lineage as the reference. This indicates that on average, genomic synteny is under weak conservation within *B. edulis*.

#### 2.9 Characterizing the super pan-genome of *B. edulis*

We found that *B. edulis* possess one of the most diverse super pan-genomes among eukaryotes. Across 200 samples with a minimum BUSCO recovery of 90% (min = 92%, med = 96.7) we identified 30,719 total orthologous gene clusters (Fig. 3B). Of these orthologous groups, 2,120 (6.9% of total) were singletons (found exclusively in one genome), 10,399 (33.9%) were rare (found in at least two genomes, but less than five), 12,597 (41%) were accessory (found in at least five genomes, but less than 95% of all genomes), and 7,256 (23.6%) were core (found in at least 95% of genomes). While the exact proportion of core and accessory orthologous groups is difficult to compare across studies, particularly among "super pan-genome" surveys of multiple taxa or species groups, *B. edulis* appears to possess among the lowest proportion of core proteins so far identified among eukaryotes. For example, two recent genomic surveys of four closely related rice (*Oryza*) and three tomato (*Solanum*) taxa found 42.6% (21,888 of 51,359 total orthologus groups), and 54% (23,839 of 40,457) core proteins respectively [32, 50]. Strikingly, *B. edulis* contains nearly half as many core proteins (as a proportion of total) as the rice super pan-genome despite possessing an estimated most recent common ancestor at least twice as old (MRCA of *B. edulis* = 2.6 MYA, MRCA of taxa in Shang et al. (2022) = 600KYA [53]). Even within an individual lineage, six out of seven lineages in *B. edulis* contain a lower percentage of core proteins (mean = 44.5%, Fig. 3C) than has been previously identified in other Fungi [34, 52]. Surprisingly, only four orthologous gene groups were lineage specific and present in at least of 90% samples within that lineage (one group in BC lineage and three in GU). These gene groups contain a predicted triacylglycerol lipase (OG0012300 in BC, EC 3.1.1.3) potentially involved in abiotic stress [22], and a predicted Caspase (OG0015485 in GU, MEROPS C14) potentially involved in apoptosis [19] or pathogen immune response [14].

#### 2.10 Identifying whether lineages of *B. edulis* differ in ecologically relevant gene families

To determine whether lineages with signatures of contemporary hybridization and/or introgression are more ecologically similar we characterized the abundance and composition of three groups of ecologically important gene families: 1) secreted and non-secreted carbohydrate active enzymes (CAZYmes) and peptidases (MEROPS) used for biodegredation and substrate modification (termed biodegredation repertoire), 2) small secreted proteins (SSPs) and membrane-bound nutrient transporters which are widely implicated in host associations (termed host-association repertoire), and 3) specific gene families involved in abiotic stress (e.g. osmotic stress, starvation stress).

##### 2.10.1 Identification of the Biodegredation repretoire of *B. edulis*

Within the biodegredation repretoire, we identified 262 CAZymes on average (112 secreted CAZymes on average) in *B. edulis* genomes, which is substantially fewer than has been found within other Boletales EMF taxa [61, 47] (Fig. 4A, Table S13-S15). These CAZymes were distributed across 112 enzyme families, of which 55 families are secreted. We also identified further reductions in enzymes targeting cellulose/hemicellulose, such as AA3 oxidoreductases (average = 9.6), and GH5 hemicellulases (average = 14.0). However, *B. edulis* appears to possess a greater number of peptidases on average (mean = 503) compared to other EMF taxa [35]. Within *B. edulis*, we found significant differences in the abundance of both CAZymes and peptidases among lineages. The GU lineage showed consistent contractions in most degradation enzyme families and possessed the fewest CAZymes and peptidases in total ($P < 0.01$, F=4.311, DF=6, one-way ANOVA, Fig. S5, Fig. S6). Several specific families were found to be enriched or depleted in specific lineages. For example, the AK lineage exhibits a unique contraction of the AA1_1 laccase family (AK average = 2.5, rest of *B. edulis* = 6.9). To determine whether the lineages differ in their overall biodegredation profiles we used PCoA to cluster specimens based on their composition and abundance of CAZymes and peptidases (Fig. 4B). We found that the AK-EU-EC lineages comprise a single overlapping cluster while all other lineages exhibit little to no overlap with any other lineage.

### 4 Conflicts of Interest

The authors declare no conflict of interest, financial or otherwise.

*NextDenovo* (Mar. 2023).

Ødum, Marius Thrane et al. (July 2024). “DeepLoc 2.1: multi-label membrane protein type prediction using protein language models”. In: *Nucleic Acids Research* 52.W1, W215–W220. ISSN: 0305-1048.

Ou, Shujun et al. (Dec. 2019). “Benchmarking transposable element annotation methods for creation of a streamlined, comprehensive pipeline”. In: *Genome Biology* 20.1, p. 275. ISSN: 1474-760X.

Palmer, Jon (Oct. 2022). *nextgenusfs/funannotate*.

Plassard, Claude et al. (2019). “Phosphorus transport in mycorrhiza: how far are we?” In: *Trends in Plant Science* 24.9, pp. 794–801.

Plett, Jonathan M et al. (Mar. 2023). “Speciation Underpinned by Unexpected Molecular Diversity in the Mycorrhizal Fungal Genus Pisolithus”. In: *Molecular Biology and Evolution* 40.3, msad045. ISSN: 1537-1719.

Rawlings, Neil D et al. (Jan. 2018). “The MEROPS database of proteolytic enzymes, their substrates and inhibitors in 2017 and a comparison with peptidases in the PANTHER database”. In: *Nucleic Acids Research* 46.D1, pp. D624–D632. ISSN: 0305-1048.

Saier Jr, Milton H et al. (Jan. 2021). “The Transporter Classification Database (TCDB): 2021 update”. In: *Nucleic Acids Research* 49.D1, pp. D461–D467. ISSN: 0305-1048.

Shang, Lianguang et al. (Oct. 2022). “A super pan-genomic landscape of rice”. en. In: *Cell Research* 32.10, pp. 878–896. ISSN: 1748-7838.

Shen, Wei et al. (Oct. 2016). “SeqKit: A Cross-Platform and Ultrafast Toolkit for FASTA/Q File Manipulation”. en. In: *PLOS ONE* 11.10, e0163962. ISSN: 1932-6203.

Steenwyk, Jacob L. et al. (Sept. 2024). “Evolutionary origin and population diversity of a cryptic hybrid pathogen”. en. In: *Nature Communications* 15.1, p. 8412. ISSN: 2041-1723.

Stein, Joshua C. et al. (Feb. 2018). “Genomes of 13 domesticated and wild rice relatives highlight genetic conservation, turnover and innovation across the genus Oryza”. en. In: *Nature Genetics* 50.2, pp. 285–296. ISSN: 1546-1718.

Tremble, Keaton et al. (Oct. 2022). “Contrasting continental patterns of adaptive population divergence in the holarctic ectomycorrhizal fungus Boletus edulis”. en. In: *New Phytologist* n/a.n/a. ISSN: 1469-8137.

Tremble, Keaton et al. (May 2024). “A revised phylogeny of Boletaceae using whole genome sequences”. EN. In: *Mycologia*. ISSN: 0027-5514.

Walker, Bruce J. et al. (Nov. 2014). “Pilon: An Integrated Tool for Comprehensive Microbial Variant Detection and Genome Assembly Improvement”. en. In: *PLOS ONE* 9.11, e112963. ISSN: 1932-6203.

Wang, Q.B. et al. (June 2005). “Boletus reticuloceps, a new combination for Aureoboletus reticuloceps”. In: *Sydowia* 57, pp. 131–136.

Wang, Xinfeng et al. (Dec. 2022). “Extensive gene flow in secondary sympatry after allopatric speciation”. In: *National Science Review* 9.12, nwac280. ISSN: 2095-5138.

Waterhouse, Robert M. et al. (Mar. 2018). “BUSCO Applications from Quality Assessments to Gene Prediction and Phylogenomics”. en. In: *Molecular Biology and Evolution* 35.3, pp. 543–548. ISSN: 0737-4038.

Williams, Matthew P et al. (Sept. 2024). “Testing times: disentangling admixture histories in recent and complex demographies using ancient DNA”. In: *Genetics* 228.1, iyae110. ISSN: 1943-2631.

Wu, Gang et al. (2022). “Evolutionary innovations through gain and loss of genes in the ectomycorrhizal Boletales”. en. In: *New Phytologist* 233.3, pp. 1383–1400. ISSN: 1469-8137.

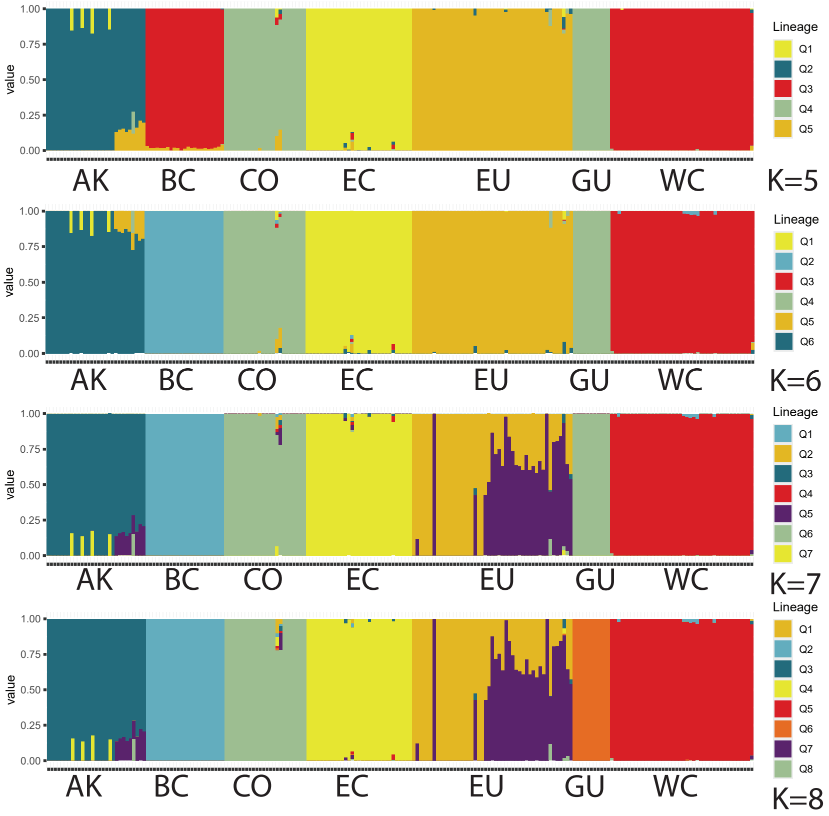


Fig. S1: Analysis of genome wide admixture using the program ADMIXTURE with values of K6 to K9.

Zhang, Han et al. (July 2018). “dbCAN2: a meta server for automated carbohydrate-active enzyme annotation”. In: *Nucleic Acids Research* 46.W1, W95–W101. ISSN: 0305-1048.

Zimin, Aleksey V. et al. (Nov. 2013). “The MaSuRCA genome assembler”. en. In: *Bioinformatics* 29.21, pp. 2669–2677. ISSN: 1367-4803.

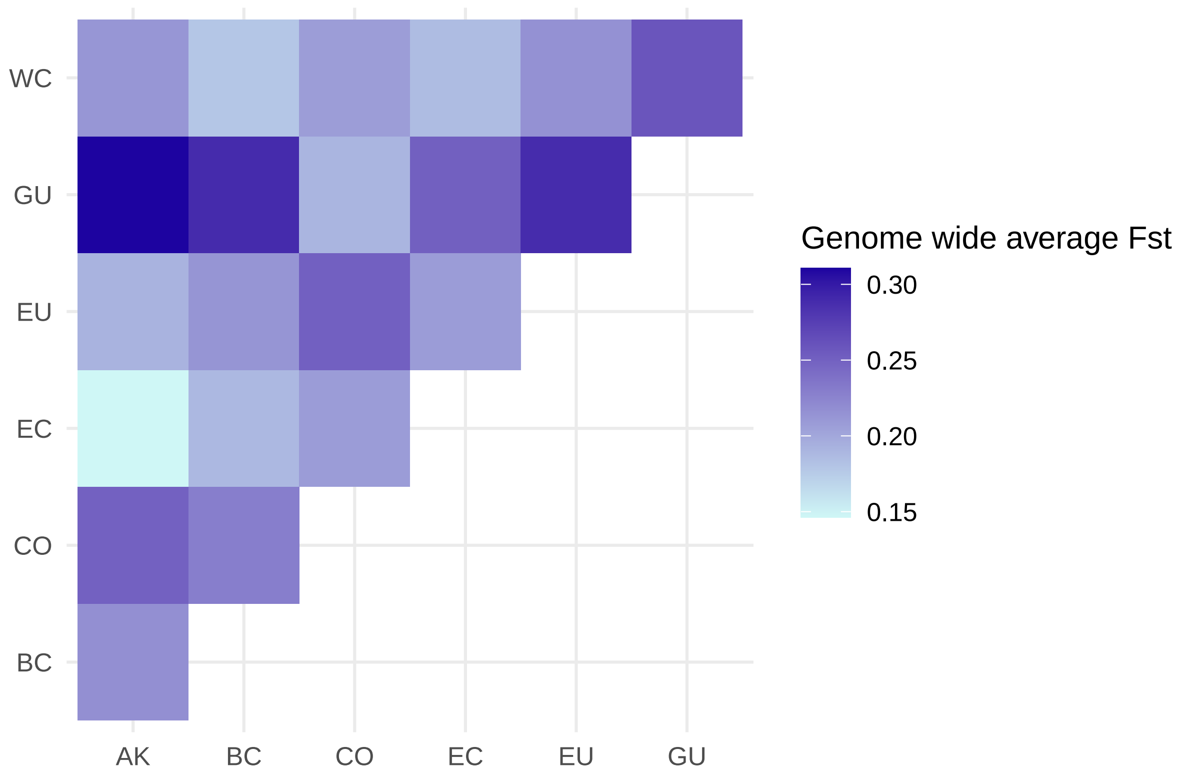


Fig. S2: Pairwise population differentiation (Fst) calculated using SNPs called against the EU chromosomal reference genome.

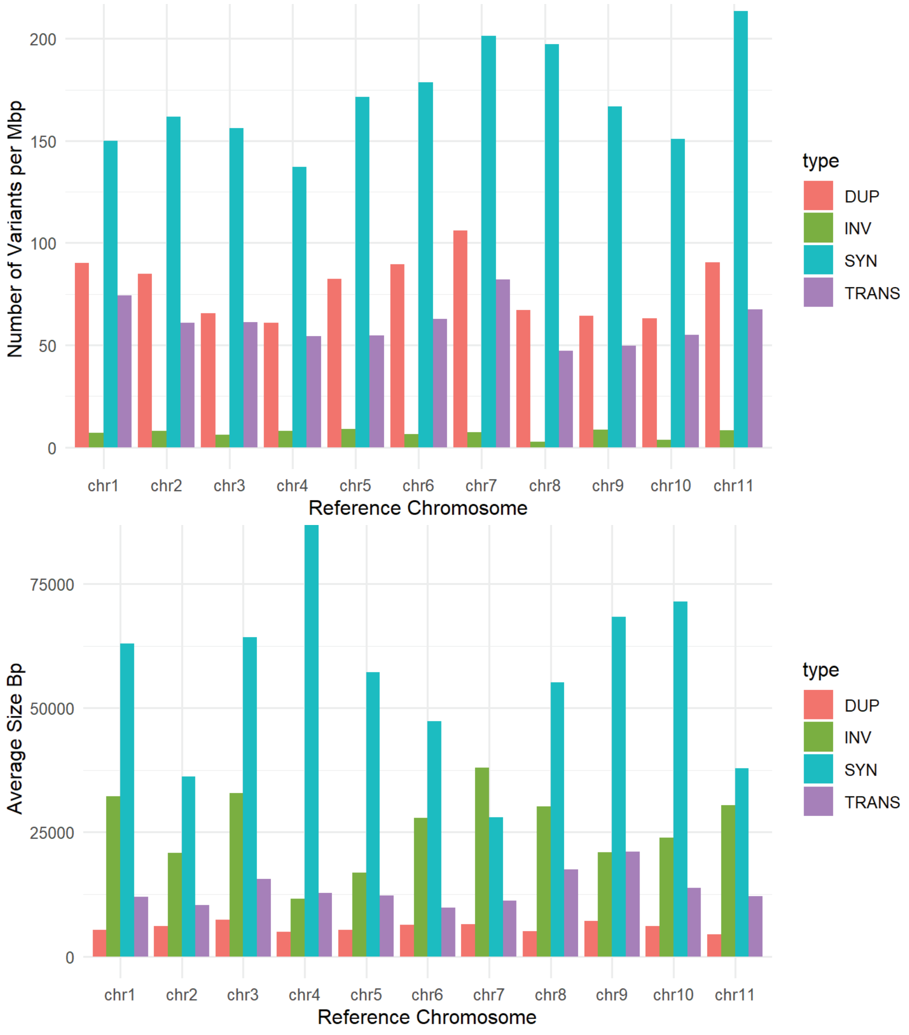


Fig. S3: The distribution of structural variants and their sizes per chromosome of the European Reference genome. Variant labels are as follows; DUP = Duplications, INV = Inversions, SYN = Syntenic blocks, TRANS = Translocations. Top shows variant counts per megabase of chromosome (Mbp), while bottom shows average variant size per chromosome.

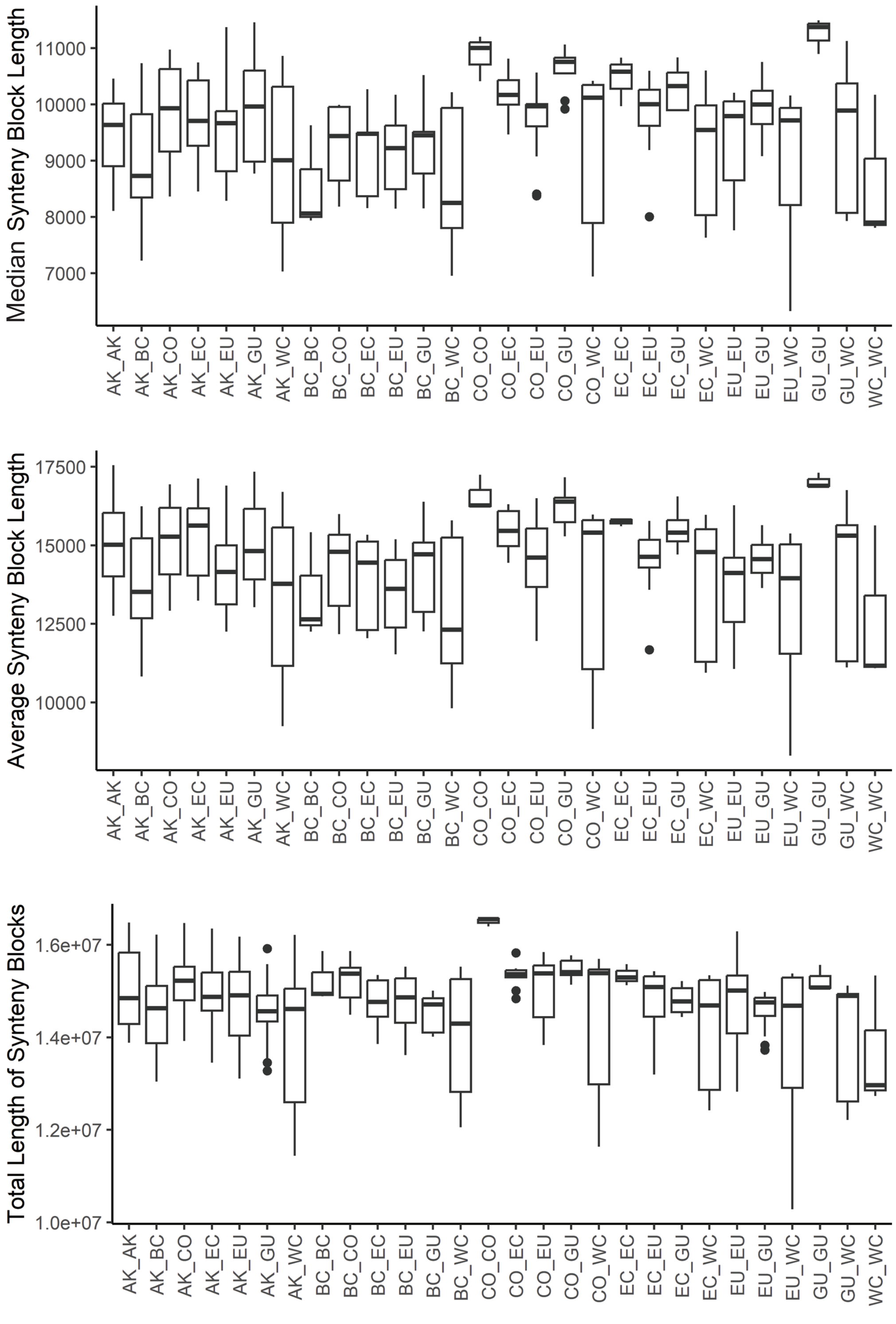


Fig. S4: Estimation of synteny blocks using orthologous gene mapping among linenage pairs.

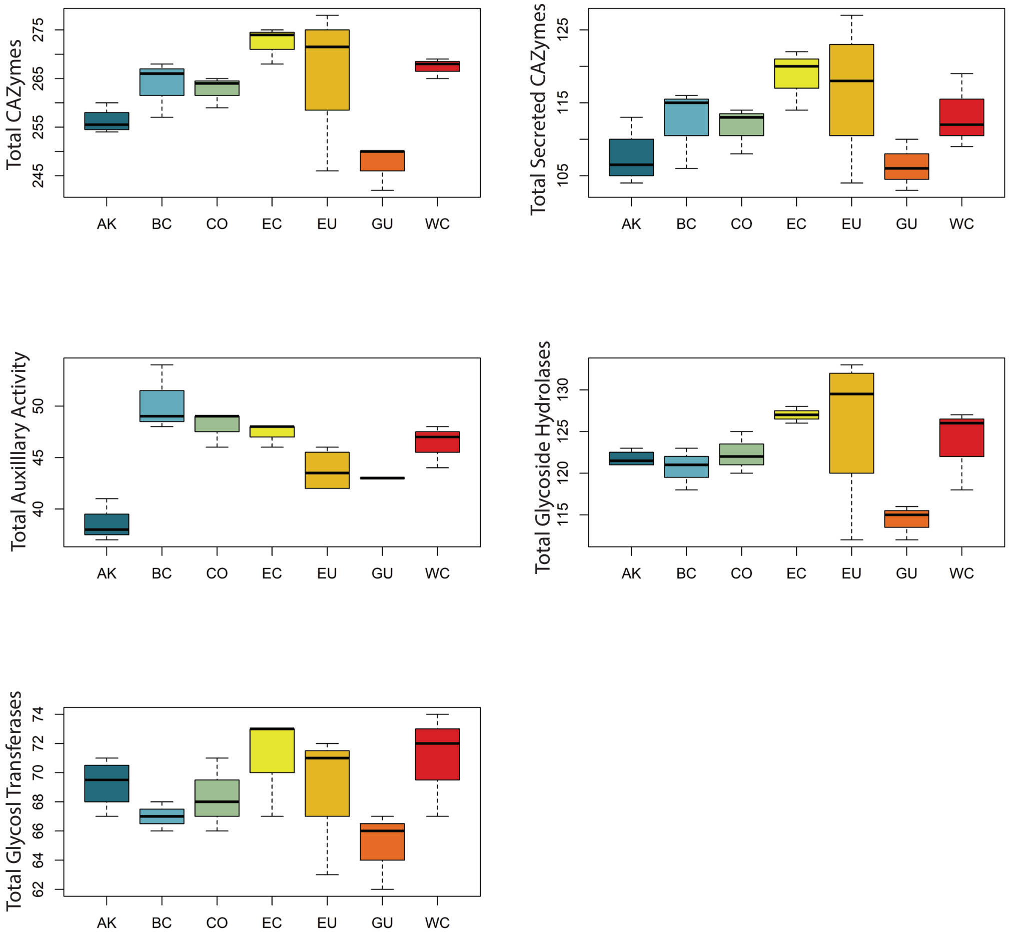


Fig. S5: Average abundance of CAZyme super-families within lineages of *B. edulis*

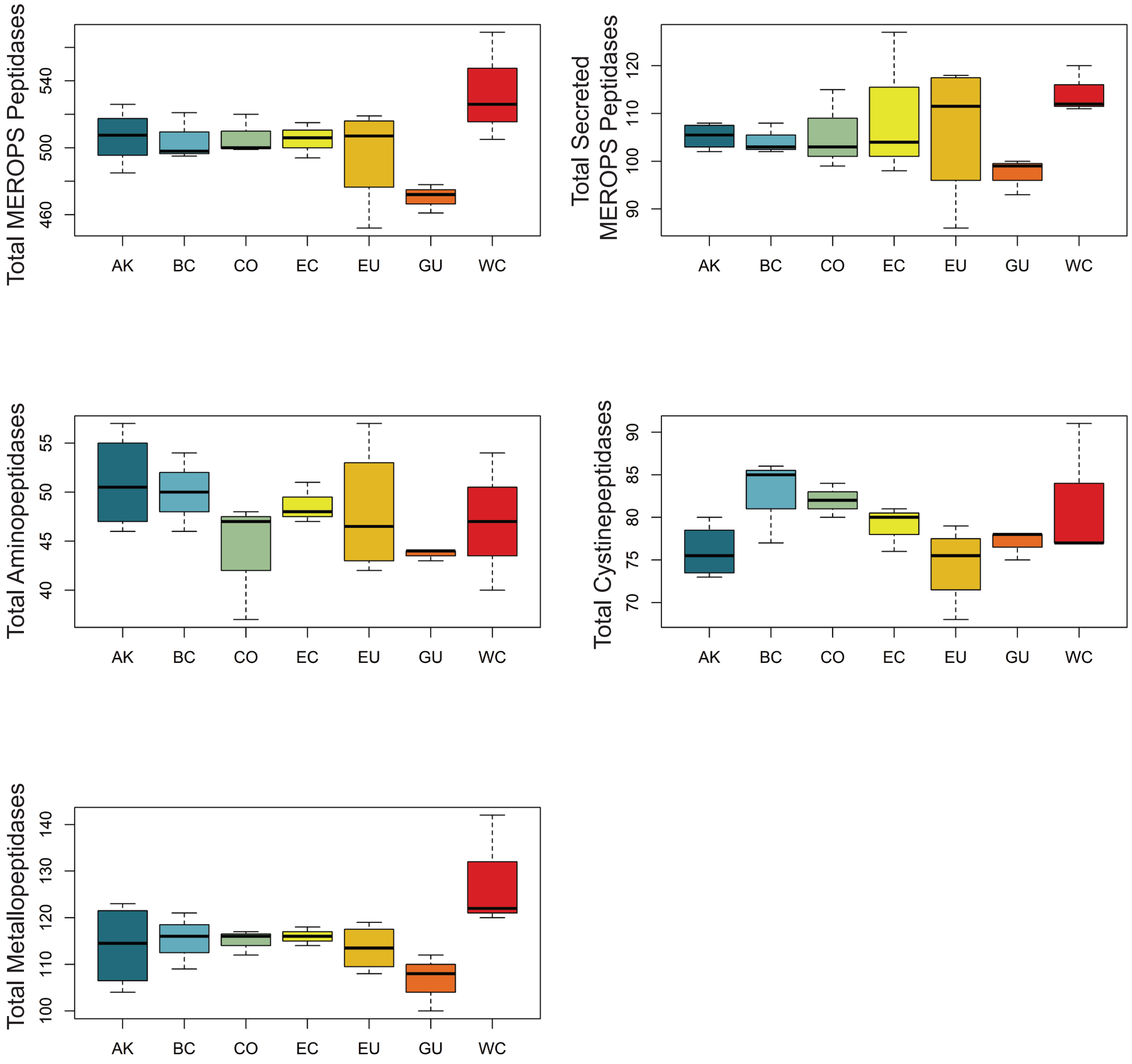


Fig. S6: Average abundance of peptidases super-families within lineages of *B. edulis*

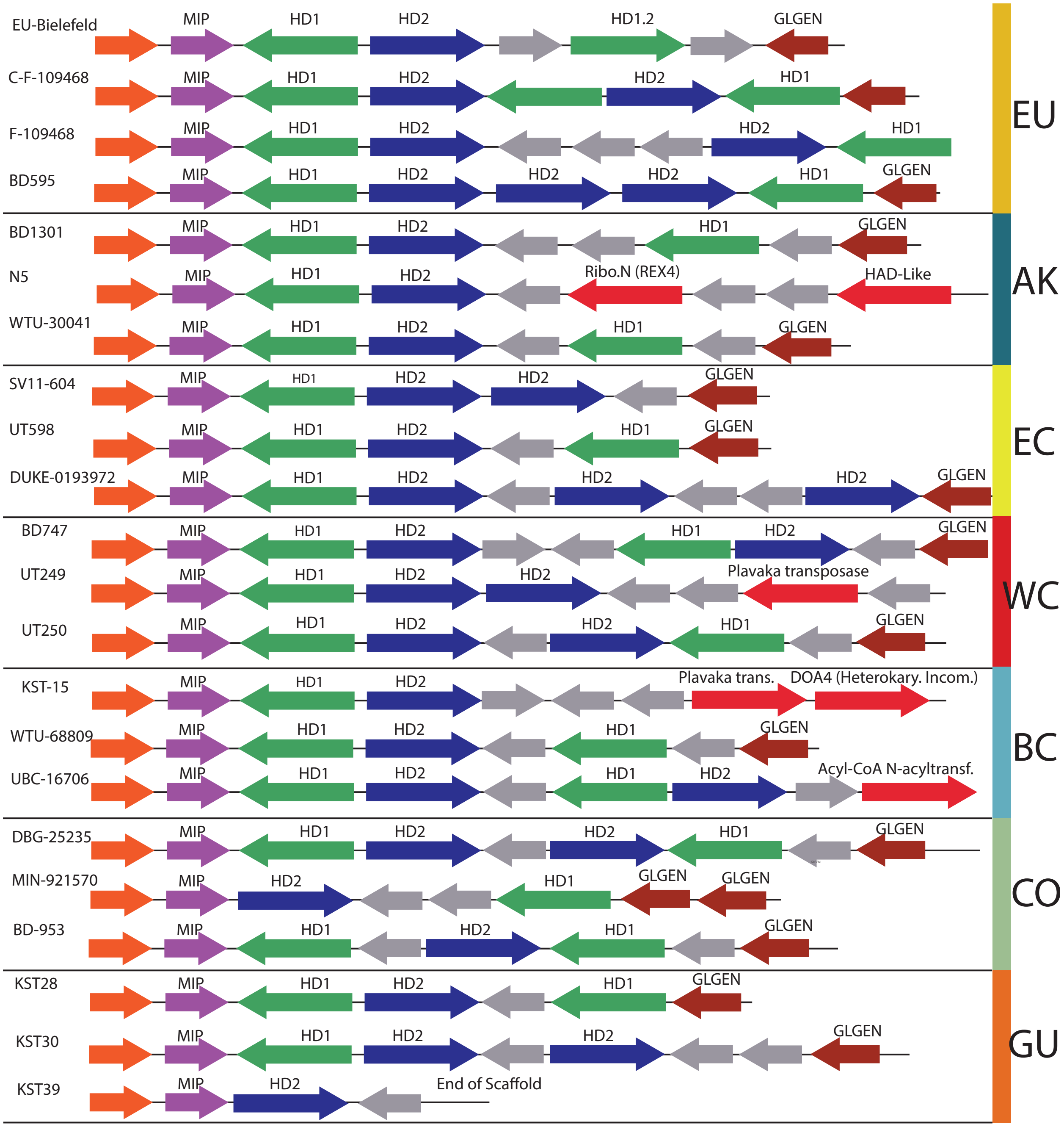


Fig. S7: Structure and orientation of the MATa homeodomain locus of *B. edulis* within 23 reference genomes of *B. edulis*. Gene blocks of the same color are orthologous with the exception of red blocks which are vaired genes with predicted function. Grey blocks are novel hypothetical proteins. MIP is an acronym for mitochondrial intermediate peptidase, HD1 stands for homeodomain locus 1, HD2 stands for homeodomain locus 2.

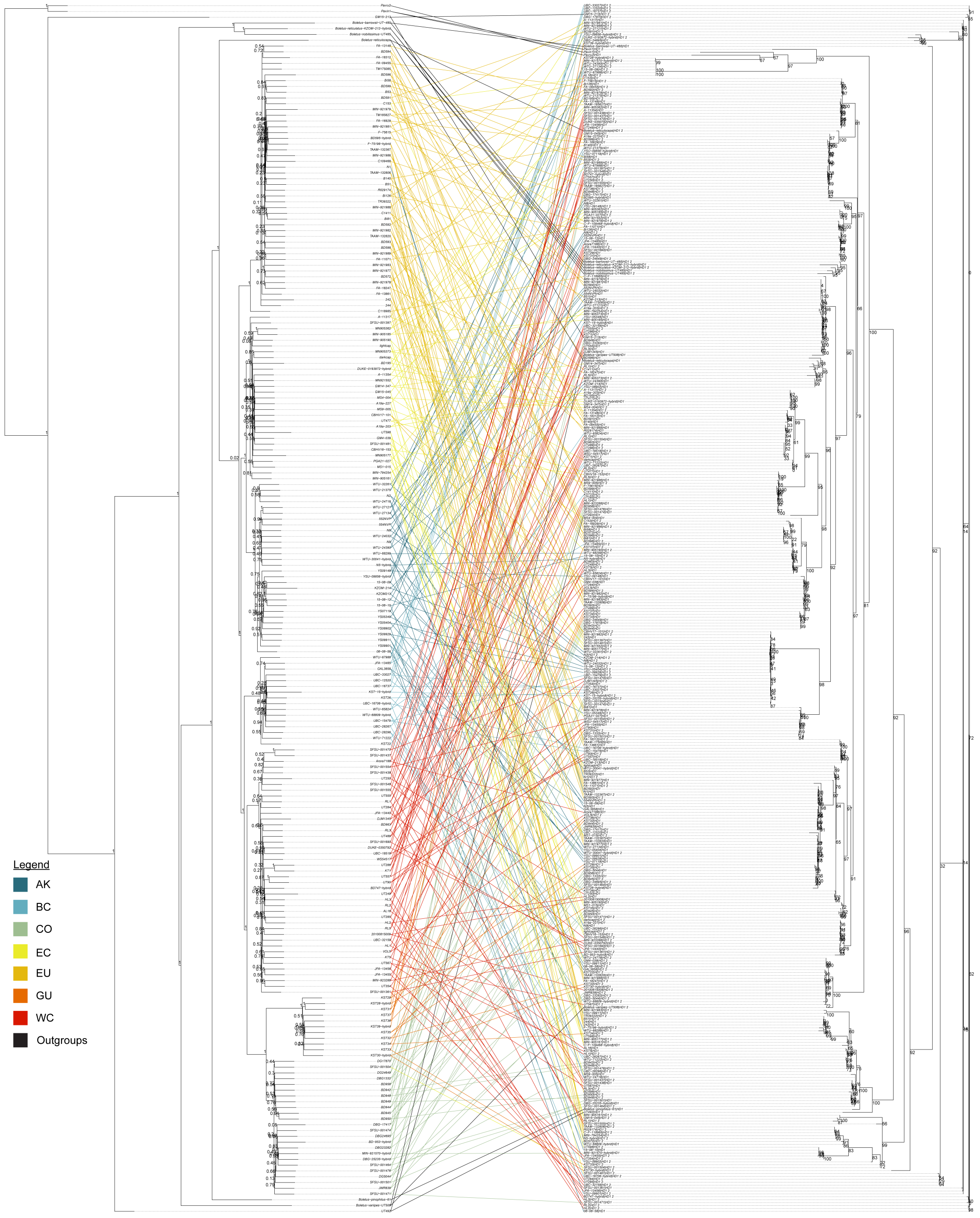


Fig. S8: Co-phylogeny of HD1 gene from MATa locus and nuclear species tree, showing lack of lineage specific nucleotide divergence of HD1.

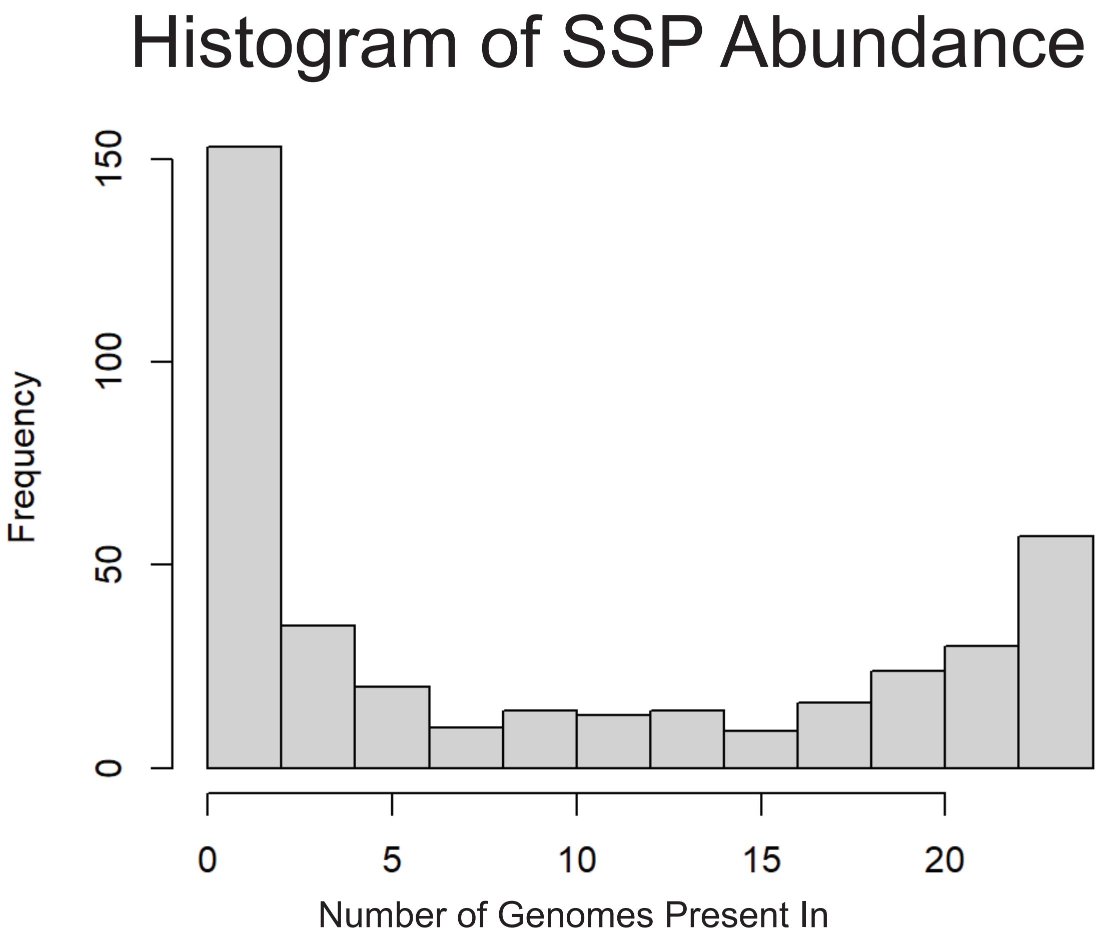


Fig. S9: Histogram of SSP abundance across the 23 reference genomes of *B. edulis*
